# Activation of MAP3K DLK and LZK in Purkinje Cells Causes Rapid and Slow Degeneration Depending on Signaling Strength

**DOI:** 10.1101/2020.10.09.332940

**Authors:** Yunbo Li, Erin M Ritchie, Christopher L. Steinke, Cai Qi, Lizhen Chen, Binhai Zheng, Yishi Jin

## Abstract

The conserved MAP3K Dual leucine zipper kinases can activate JNK via MKK4 or MKK7. Vertebrate DLK and LZK share similar biochemical activities and undergo auto-activation upon increased expression. Depending on cell-type and nature of insults DLK and LZK can induce pro-regenerative, pro-apoptotic or pro-degenerative responses, although the mechanistic basis of their action is not well understood. Here, we investigated these two MAP3Ks in cerebellar Purkinje cells using loss- and gain-of function mouse models. While loss of each or both kinases does not cause discernible defects in Purkinje cells, activating DLK causes rapid death and activating LZK leads to slow degeneration. Each kinase induces JNK activation and caspase-mediated apoptosis independent of each other. Significantly, deleting CELF2, which regulates alternative splicing of *Mkk7*, strongly attenuates Purkinje cell degeneration induced by activation of LZK, but not DLK. Thus, controlling the activity levels of DLK and LZK is critical for neuronal survival and health.

## Introduction

Mitogen-activated protein kinase (MAPK) signaling pathways play important roles in neuronal development and function, and aberrant regulation of MAP kinases is associated with many neurological diseases, such as Parkinson’s disease (PD), amyotrophic lateral sclerosis (ALS) and Alzheimer’s disease (AD) (Thomas and Huganir, 2004; Schellino et al., 2019; Hotamisligil and Davis, 2016; Hollville et al., 2019; Adib et al., 2018). The MAPK cascade involves MAP3Ks (MAP kinase kinase kinases), MAP2Ks and MAPKs that together form a phosphorylation relay and activate downstream signaling events in response to external or internal stimuli. The mammalian MAP3K DLK (Dual leucine zipper kinase, or MAP3K12) and LZK (Leucine zipper kinase, or MAP3K13) are members of an evolutionarily conserved family that includes *C. elegans* DLK-1 and Drosophila DLK/Wallenda (Jin and Zheng, 2019). These MAP3Ks act as upstream kinases for JNK and p38 MAP kinase, and are now known as key players in neuronal stress response network both under acute injury and in chronic neurodegenerative diseases (Jin and Zheng, 2019; Adib et al., 2018; Farley and Watkins, 2018). An emerging theme is that while activation of these kinases triggers seemingly common pathways, the outcome is highly context-specific both in terms of cell types and forms of insults.

Both DLK and LZK show broad expression in the nervous system. Several studies have investigated roles of DLK in the development of the nervous system. Constitutive *Dlk* knockout mice die perinatally, and different regions of developing brain display varying degrees of altered axon fibers, abnormal synapses and increased neuronal survival (Hirai et al., 2006; Hirai et al., 2011; Nakata et al., 2005; Collins et al., 2006; Lewcock et al., 2007). However, mice with adult deletion of *Dlk* survive and show no detectable abnormalities (Le Pichon et al., 2017; Tedeschi and Bradke, 2013). Under traumatic insults, DLK activity is reported to increase and trigger a variety of cellular responses. For example, sciatic nerve injury induces DLK-dependent pro-regenerative responses in dorsal root ganglia (DRG) sensory neurons (Shin et al., 2012; Shin et al., 2019). In the central nervous system (CNS), optic nerve injury up-regulates DLK expression in retinal ganglion cells (RGC), which triggers cell death in many RGCs and also promotes axon growth from surviving RGCs (Watkins et al., 2013; Welsbie et al., 2013). In a mouse model for stroke, increased DLK expression in pre-motor cortex is suggested to promote motor recovery (Joy et al., 2019). Increased DLK activity is also reported in animal models of neurodegeneration, and genetically or pharmacologically inhibiting DLK in the aged PS2APP mice for AD and the SOD1^G93A^ mice for ALS has resulted in some neuroprotective effects (Chen et al., 2008; Le Pichon et al., 2017). Intriguingly, in human iPSC derived neurons treated with ApoE4, a protein associated with an increased risk for AD, DLK is rapidly up-regulated and enhances transcription of APP (Huang et al., 2017). As numerous approaches now target DLK for drug discovery (Siu et al., 2018), it is important to investigate how the pleiotropic effects of manipulating DLK in different cell types influence disease progression. In comparison, despite the fact that LZK was also discovered 20 years ago (Sakuma et al., 1997; Holzman et al., 1994), the *in vivo* roles of LZK are only beginning to be explored. In a mouse model of spinal cord injury, LZK is upregulated in astrocytes and mediates reactive astrogliosis (Chen et al., 2018). Emerging studies show that LZK can cooperate with DLK in RGC to promote cell death after optic nerve injury and in DRG for axon degeneration (Welsbie et al., 2017; Summers et al., 2020).

Here, we dissect the roles of the two kinases in the cerebellar Purkinje cells. We analyzed genetic deletion mice for each kinase, and also developed transgenic mice that allow for Cre-mediated expression of DLK or LZK. Biochemical studies have shown that DLK and LZK undergo auto-activation via leucine-zipper mediated dimerization, and such auto-activation is dependent on the protein abundance (Nihalani et al., 2000; Ikeda et al., 2001b). Therefore, elevating expression of DLK or LZK is a proxy to its activation of the downstream signal transduction. We find that deletion of DLK and/or LZK, singly or in combination, from Purkinje cells, does not affect their development and postnatal growth. In contrast, induced expression of DLK in Purkinje cells causes rapid degeneration, whereas elevating LZK expression induces a slow degeneration. Strikingly, we find that deleting the RNA splicing factor CELF2 ameliorates Purkinje cell degeneration induced by LZK, but not DLK, partly via regulating alternative splicing of *Mkk7*, a MAP2K. These findings provide important insights to the understanding of neurodegenerative processes.

## Results

### Normal development of cerebellar Purkinje cells in the absence of DLK and LZK

Both DLK and LZK are expressed in cerebellar neurons, with high levels of DLK observed in the molecular layer of adult cerebellum (Hirai et al., 2005; Suenaga et al., 2006; Goodwani et al., 2020). *Dlk* knockout (KO) mice die soon after birth, and cerebellar architecture is grossly normal (Hirai et al., 2006). The roles of *Lzk* in neuronal development remain unknown. To address the function of *Lzk* and further probe into the interactions between the two kinases, we generated an *Lzk* KO mouse line using CRISPR-editing to delete the entire kinase domain (Figure 1-figure supplement 1 A-B), which also resulted in a frameshift and produced no detectable LZK proteins by immunoprecipitation and western blotting analysis (Figure 1-figure supplement 1 C). The *Lzk* KO mice are viable, and grow indistinguishably from littermates under standard housing conditions. Histological analysis of cerebellar tissue sections using hematoxylin and eosin staining revealed no discernible defects in overall cellular architecture in two-month old (P60) *Lzk* KO mice (Figure 1 A). Immunostaining using antibodies to Calbindin, which specifically labels Purkinje cells, showed that the position, number and gross morphology of Purkinje cells were comparable between *Lzk* KO and control (Figure 1 B-D). The molecular layer thickness, which is a sensitive assessment for disruption of the dendrites of Purkinje cells (White et al., 2014; Hansen et al., 2013; White et al., 2016; White and Sillitoe, 2017), was also normal (Figure 1 E).

**Figure 1.**
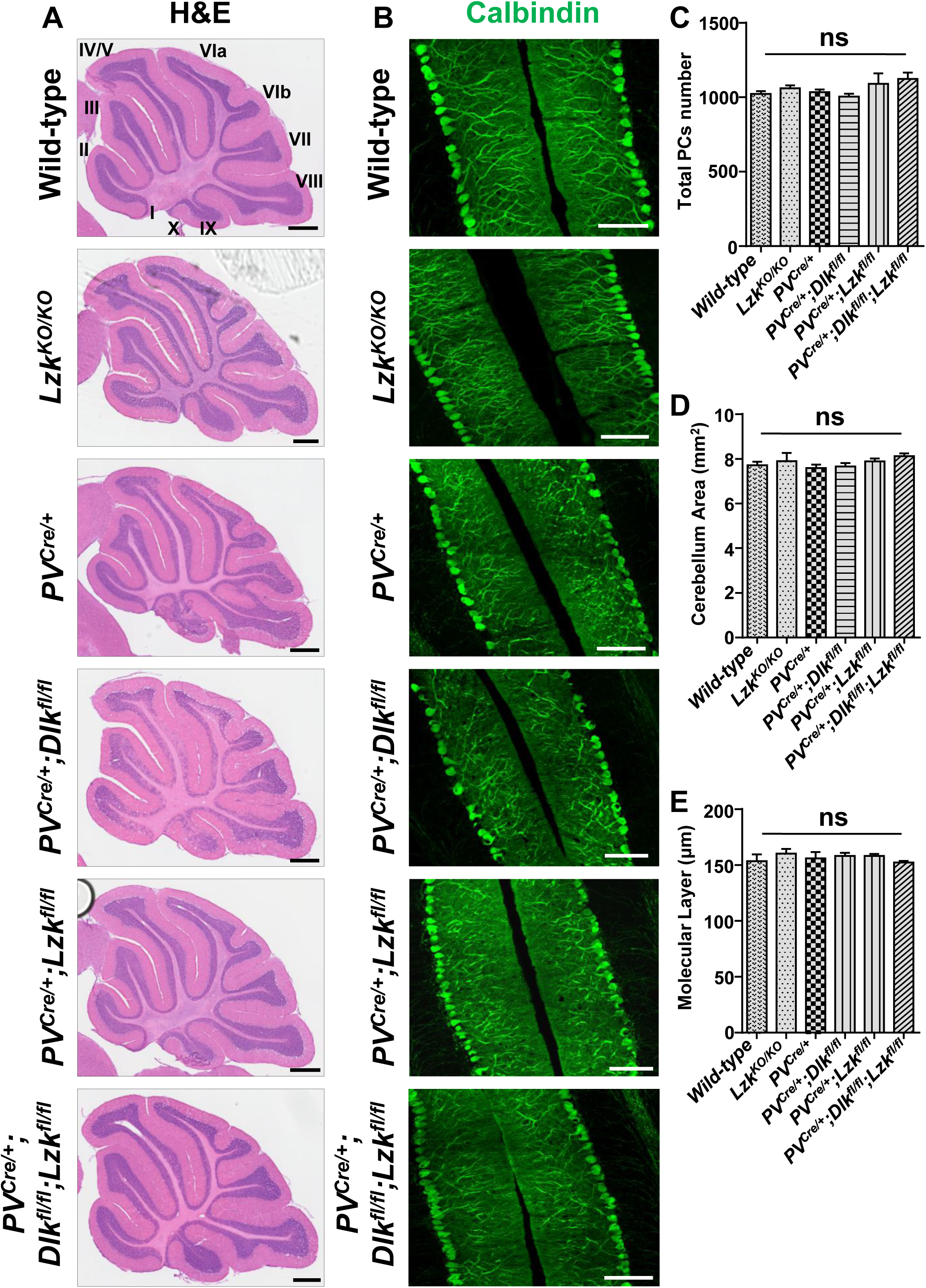
Morphology of cerebellar Purkinje cells is normal in the absence of LZK and DLK. A. Representative images of hematoxylin and eosin staining of cerebellar sections from P60 mice of genotypes indicated. Scale bars: 500 μm. B. Representative images of Calbindin staining of cerebellar sections from P60 mice of genotypes indicated. Scale bars: 100 μm. C. Quantification of total number of Purkinje cells in all cerebellar lobules. D. Quantification of cerebellum area, with perimeters measured by outlining the outer edge of midline sagittal sections of the cerebella. E. Quantification of the molecular layer thickness in cerebellar lobules V-VI. (C-E). n ≥ 3 animals per genotype, and 3 sections/animal; data shown as means ± SEM. Statistics: One-way ANOVA; ns. no significant.

It has recently been shown that *Dlk* and *Lzk* can act synergistically in injured RGC or DRG neurons (Welsbie et al., 2017; Summers et al., 2020). We therefore tested if loss of both DLK and LZK might affect cerebellar neurons. We bred floxed (*fl*) *Dlk* or *Lzk* KO mice to a parvalbumin-Cre driver line (*PV*^*Cre*^) (Hippenmeyer et al., 2005), and obtained *PV*^*Cre*/+^;*Dlk*^*fl/fl*^;*Lzk*^*fl/fl*^ mice, along with *PV*^*Cre*/+^;*Dlk*^*fl/fl*^ and *PV*^*Cre*/+^;*Lzk*^*fl/fl*^ (Figure 1-figure supplement 1 D-G). Cre recombinase from *PV*^*Cre*^ line is active in Purkinje cells as early as P4 (Hippenmeyer et al., 2005). We detected reduced protein levels of DLK and LZK in cerebellar extracts for *PV*^*Cre*/+^;*Dlk*^*fl/fl*^;*Lzk*^*fl/fl*^, *PV*^*Cre*/+^;*Dlk*^*fl/fl*^ and *PV*^*Cre*/+^;*Lzk*^*fl/fl*^ mice, respectively (Figure 1-figure supplement 1 C, H). The overall cerebellar tissue organization, revealed by hematoxylin and eosin staining, was indistinguishable among test and control mice of P60 age (Figure 1 A). Calbindin immunostaining showed that the total number of Purkinje cells and the molecular layer thickness of cerebellum were comparable in single or double gene deletion of each kinase (Figure 1 B-E). Additionally, GFAP immunostaining for cerebellar astrocytes revealed no detectable difference among different genotypes of mice (Figure 1-figure supplement 1 I-J). All mutant mice also showed postnatal growth, measured by body weight, comparable to the control mice under same housing conditions (Figure 1-figure supplement 1 K). These data show that DLK and LZK are not required for the postnatal development of Purkinje cells.

### Elevating DLK expression in Purkinje cells causes rapid degeneration via apoptosis

Increased expression of DLK or LZK has been reported under traumatic injury or other stress conditions (Shin et al., 2012; Shin et al., 2019; Watkins et al., 2013; Welsbie et al., 2013; Joy et al., 2019; Chen et al., 2018; Huang et al., 2017). We next investigated how elevating expression of DLK and LZK, hence activation of these kinases, affects neurons. To this end, we generated two transgenic mouse lines by inserting a Cre-inducible transgene of *Dlk* or *Lzk* at the *Hipp11* (*H11*) locus, named *H11*-*Dlk*^*iOE*^ or *H11*-*Lzk*^*iOE*^, respectively (Figure 2-figure supplement 1 A-B). In each transgene, the induced expression can be readily assessed by a tdTomato reporter fused in-frame to the C-terminus of DLK or LZK through the T2A self-cleaving peptides. By RNA-seq analysis we detected comparable levels of tdTomato mRNAs produced from each transgene following expression of Cre recombinase (Figure 2-figure supplement 1 C). After outcrossing to C57BL/6J background, these mice were bred to the *PV*^*Cre*^ line. In *PV*^*Cre*/+^;*Dlk*^*iOE*/+^ or *PV*^*Cre*/+^;*Lzk*^*iOE*/+^ heterozygous mice, we observed tdTomato reporter expression correlating with the timing of parvalbumin expression (Figure 2-figure supplement 1 D), and increased DLK or LZK expression was detected in cerebellar lysates (Figure 2-figure supplement 1 E-H).

Elevating DLK expression in *PV*^*Cre*/+^;*Dlk*^*iOE*/+^ mice caused abnormalities noticeable as early as P6. The pups were smaller than littermate controls (Figure 2 A-B), and exhibited abnormal movements (supplement video 1). These pups all died around P21. The cerebella of *PV*^*Cre*/+^;*Dlk*^*iOE*/+^ were much smaller than those of littermate controls (Figure 2 C; Figure 2-figure supplement 2 A). Histological analysis revealed grossly abnormal lobular morphology (Figure 2-figure supplement 2 B). By Calbindin immunostaining we detected a rapid degeneration of Purkinje cells, with a nearly complete (∼98%) cell loss by P21 (Figure 2 D-E). The molecular layer of the cerebellum in these mice was significantly thinner than that in the littermate control mice from P10 to P21 (Figure 2 F). Purkinje cell degeneration is known to be associated with increased reactivity of astrocytes and microglia (Cvetanovic et al., 2015; Lobsiger and Cleveland, 2007; Lattke et al., 2017). Indeed, we observed increased expression of GFAP and IBA1 (detecting both microglia and macrophage) in these mice at P21, compared to control mice (Figure 2-figure supplement 2 C-F). Some IBA1 positive microglia were closely associated with tdTomato labeled Purkinje cells (Figure 2-figure supplement 2 C), suggesting that dying cells might be phagocytosed.

**Figure 2.**
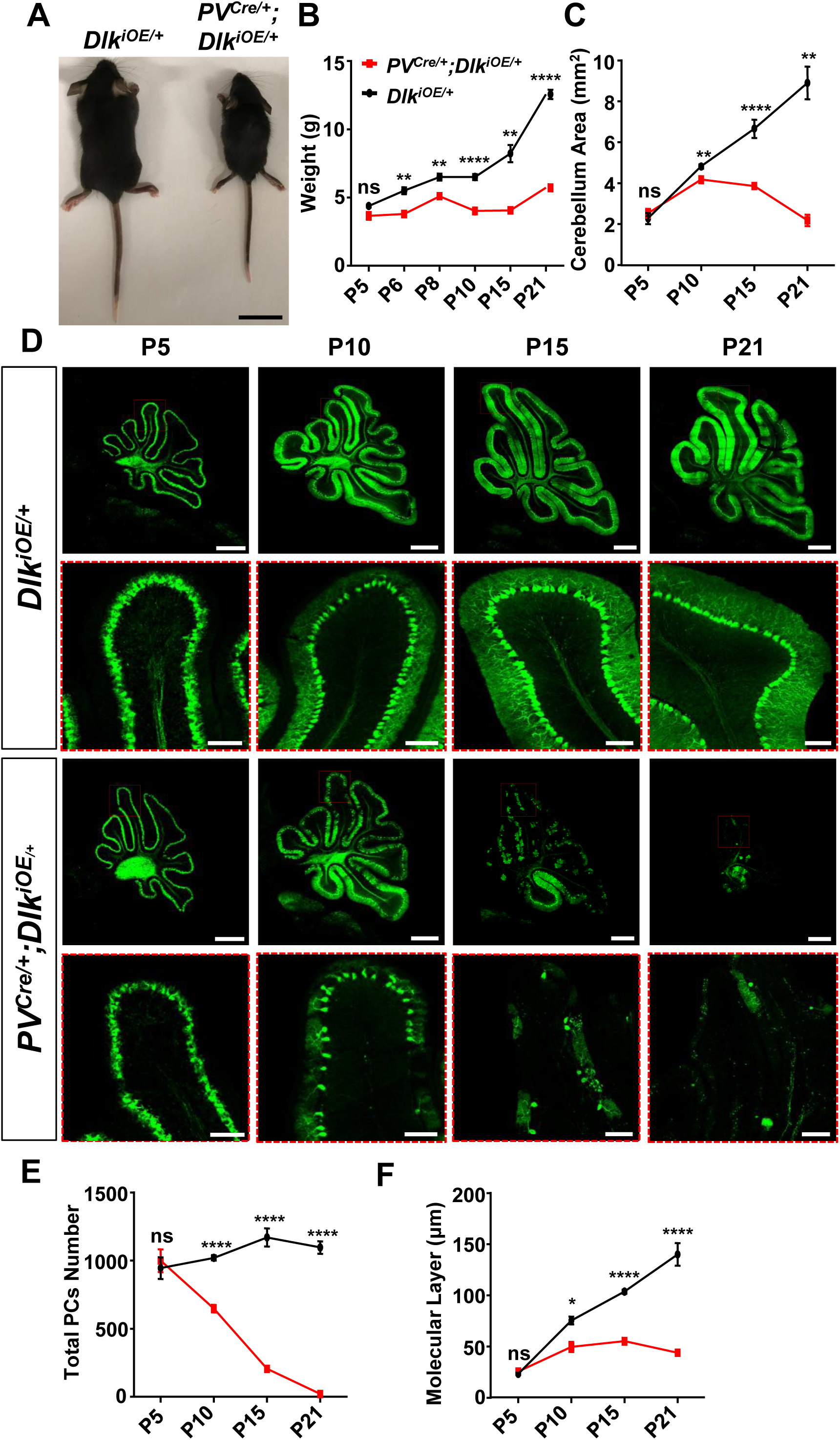
Induced expression of DLK using parvalbumin-cre (*PV*^*Cre*^) causes animal growth defects and rapid degeneration of Purkinje cells. A. Representative image of P21 pups of genotypes indicated, and pups with induced DLK expression in PV^+^ neurons are smaller than siblings. Scale bar: 2 cm. B. Quantification of the body weight from P5 to P21. n ≥ 3 per genotype for each time point. C. Quantification of the cerebellum area from P5 to P21. n ≥ 3 per genotype for each time point. D. Representative images of Calbindin staining of cerebellar sections of littermates from mating parents *PV*^*Cre*/+^ to *Dlk*^*iOE*/+^ (short for *H11-LSL-Dlk*^*iOE*^ transgene) at indicated postnatal days. Red boxes are enlarged to show that induced expression of DLK causes a total loss of Purkinje cells by P21. Scale bars: 500 μm (upper panels), 100 μm (lower panels). E. Quantification of total number of Purkinje cells in all cerebellar lobules. F. Quantification of the molecular layer thickness in cerebellar lobules V-VI. Color representation for genotypes in C, E, and F is the same as in B. (E, F). n ≥ 3 animals per genotype, 3 sections/animal; data shown as means ± SEM. Statistics for B-C, E-F: Student’s unpaired t-test; ns, no significant; *, p<0.05; **, p<0.01; ****, p<0.0001.

To determine if targeted expression of DLK induced activation of the JNK signaling, we co-immunostained for phospho-c-Jun (p-c-Jun) and Calbindin on cerebellar tissues of P6 mice. While many p-c-Jun signals were likely from granule neurons as they did not overlap with Calbindin^+^ Purkinje cells in both mutant and control mice, we observed that DLK activation induced substantially increased p-c-Jun in Purkinje cells of *PV*^*Cre*/+^;*Dlk*^*iOE*/+^ mice, compared to littermate controls (Figure 3 A-B). We also asked if the loss of Purkinje cells involved apoptosis using the TUNEL assay. During early postnatal cerebellar development, multiple types of cells undergo apoptosis, including those in the granular and the molecular layers of the cerebellar cortex (Cheng et al., 2011). Indeed, in the control littermates, we observed many TUNEL signals at P5, which decreased over the following postnatal days (Figure 3-figure supplement 1 A-B). In the *PV*^*Cre*/+^;*Dlk*^*iOE*/+^ mice, the number of apoptotic cells at P5 was comparable to that in control, but continued to rise over the next 10 days, reaching peak levels around P15 (Figure 3-figure supplement 1 A-B). Importantly, some TUNEL signals detected in P15 *PV*^*Cre*/+^;*Dlk*^*iOE*/+^ mice co-localized with tdTomato-labeled Purkinje cells (Figure 3-figure supplement 1 C). Furthermore, a significant portion of the Purkinje cells in the P15 *PV*^*Cre*/+^;*Dlk*^*iOE*/+^ mice were positively stained for cleaved (and thus activated) caspase-3 (Figure 3 C-D), a molecular marker for apoptosis (Elmore, 2007). Collectively, these results show that elevating DLK expression in Purkinje cells activates the JNK pathway and causes early-onset, rapid degeneration through apoptotic cell death.

**Figure 3.**
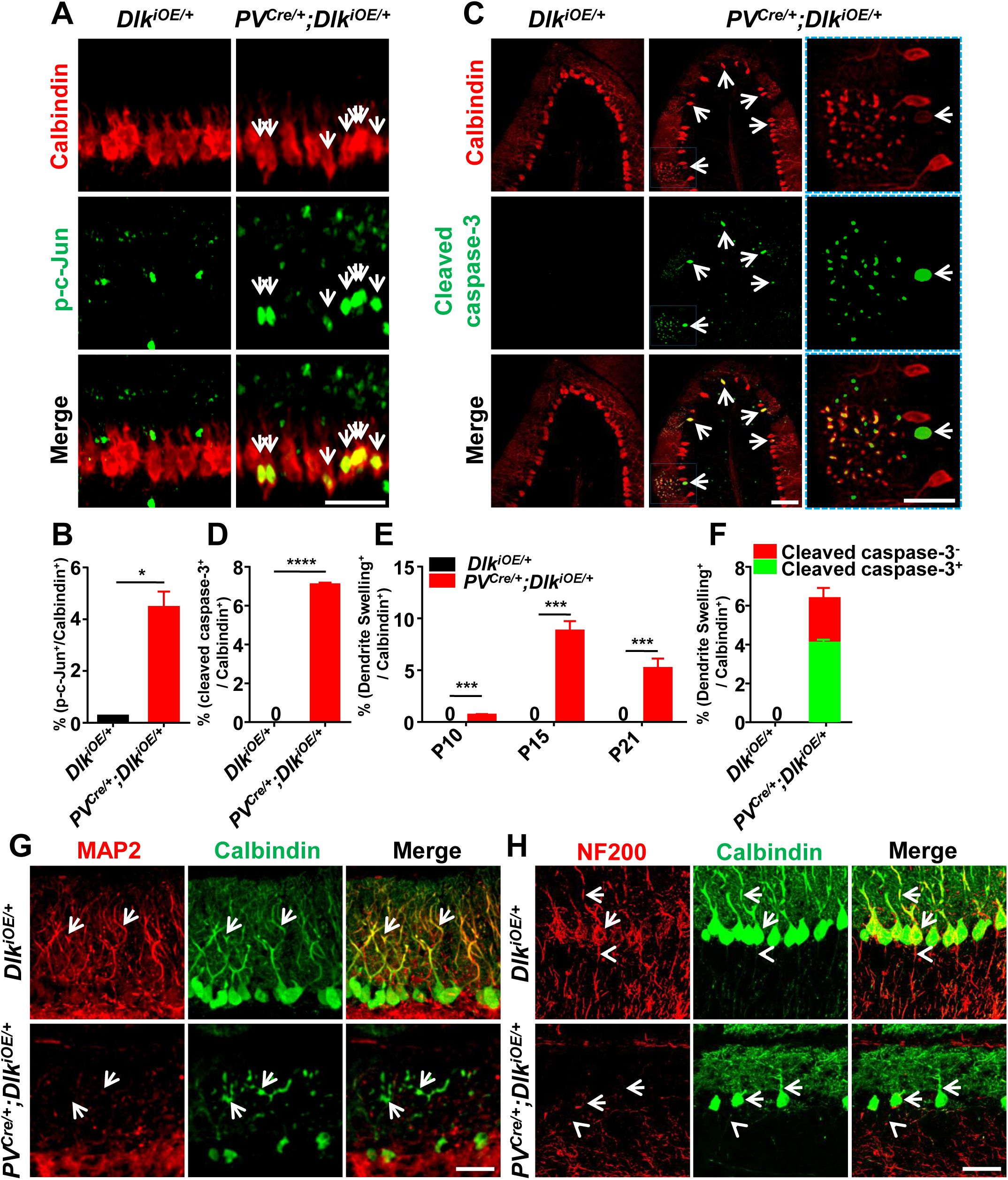
Induced expression of DLK activates JNK signaling and apoptosis, and disrupts dendritic cytoskeleton of Purkinje cells. A. Representative images of co-immunostaining of p-c-Jun and Calbindin in Purkinje cells in P6 mice of genotypes indicated. Arrows point to p-c-Jun immunostaining signals in nuclei of Purkinje cells. Scale bar: 100 μm. B. Quantification of percentage of p-c-Jun^+^ signals per total Purkinje cells in P6 mice of *Dlk*^*iOE*/+^ and *PV*^*Cre*/+^;*Dlk*^*iOE*/+^. C. Representative images of cerebellar lobule III of P15 mice, co-immunostained for Calbindin and cleaved caspase-3, with arrows and enlarged blue boxes showing cleaved caspase-3^+^ signals in the swollen dendrites of Purkinje cells of *PV*^*Cre*/+^;*Dlk*^*iOE*/+^ mice. Scale bars: 100 μm for the left and middle columns, 50 μm for the right. D. Quantification of percentage of cleaved caspase-3^+^ cells in total Purkinje cells in P15 mice of *Dlk*^*iOE*/+^ and *PV*^*Cre*/+^;*Dlk*^*iOE*/+^, and none detected in *Dlk*^*iOE*/+^. E. Quantification of percentage of Purkinje cells with swelling dendrites in mice of *Dlk*^*iOE*/+^ and *PV*^*Cre*/+^;*Dlk*^*iOE*/+^ from P10 to P21. n ≥ 3 animals per genotype at each time point, none detected in *Dlk*^*iOE*/+^. F. Quantification of percentage of Purkinje cells containing cleaved caspase-3^+^ in swelling dendrites in total Purkinje cells in P15 mice of *Dlk*^*iOE*/+^ and *PV*^*Cre*/+^;*Dlk*^*iOE*/+^. G. Representative images of Purkinje cells of P15 mice co-immunostained for MAP2 and Calbindin. Swelling dendrites in Purkinje cells of *PV*^*Cre*/+^;*Dlk*^*iOE*/+^ mice have little expression of MAP2; arrows point to dendrites with intensity difference between the two genotypes. Scale bar: 50 μm. H. Representative images of Purkinje cells of P15 mice co-immunostained for NF-200 and Calbindin, showing that induced DLK expression reduces NF-200 staining in dendrites of Purkinje cells (arrows). Scale bar: 50 μm. (B, D, E, F). n = 3 animals per genotype, minimal 3 sections/animal; data shown are means ± SEM. Statistics: Student’s unpaired t-test; ns, no significant; *, p<0.05; ***, p<0.001; ****, p<0.0001.

### DLK activation disrupts dendritic cytoskeleton

DLK is known to be localized to neuronal processes (Suenaga et al., 2006) and regulates microtubule stability (Simard-Bisson et al., 2017; Valakh et al., 2015; Hirai et al., 2011). We performed immunostaining using anti-DLK antibodies on cerebellar section of *PV*^*Cre*/+^;*Dlk*^*iOE*/+^ mice and detected DLK expression in the somas, dendrites and axons of Purkinje cells (Figure 3-figure supplement 2). In these mice, a substantial portion of Purkinje cells, as visualized by Calbindin, showed dendrite swelling (Figure 3 C, E). Moreover, ∼65% of Purkinje cells that were positively stained for cleaved caspase-3 showed dendrite swelling (Figure 3 F). The percentage of Purkinje cells with dendrite swelling was highest around P15 (Figure 3 E), consistent with the time course of Purkinje cell death caused by increased DLK expression.

Dendrite swelling is associated with major disorganization of the cytoskeleton network (Cupolillo et al., 2016; Liu et al., 2015; Hoskison et al., 2007). We next assessed how the microtubule cytoskeleton was altered by immunostaining for microtubule-associated protein 2 (MAP2), which is expressed in dendrites of Purkinje cells (Dehmelt and Halpain, 2005). We found that MAP2 levels were significantly decreased in dendrites of Purkinje cells in *PV*^*Cre*/+^;*Dlk*^*iOE*/+^ mice, compared to the levels of Calbindin as well as to control mice at P15 (Figure 3 G). The neurofilament protein NF-200 is present in both dendrites and axons and implicated in axon growth and regeneration (Wang et al., 2012). By immunostaining, we observed decreased levels of NF-200 in axons and dendrites of Purkinje cells in *PV*^*Cre*/+^;*Dlk*^*iOE*/+^ mice (Figure 3 H). Together, these data are consistent with the notion that DLK regulates the neuronal cytoskeleton, and further suggest that the dendritic cytoskeleton in Purkinje cells is highly susceptible to disruption upon aberrant activation of DLK.

### Elevating LZK expression in Purkinje cells causes progressive degeneration

In contrast to the early lethality of *PV*^*Cre*/+^;*Dlk*^*iOE*/+^ pups, the *PV*^*Cre*/+^;*Lzk*^*iOE*/+^ mice survived to older adults (observed up to 8 months). The adult mice had low body weight, compared to control mice (Figure 4-figure supplement 1 A-B). Histological analysis showed the presence of all lobular structures in P120 *PV*^*Cre*/+^;*Lzk*^*iOE*/+^ mice (Figure 4-figure supplement 1 C). Calbindin immunostaining revealed morphological abnormalities of Purkinje cells around P15, with severity and cell loss increasing from P21 to P120 (Figure 4 A-B). The area of cerebellum and the molecular layer were also reduced significantly (Figure 4 C-D). Intriguingly, we detected a stronger fluorescence intensity of tdTomato and LZK immunostaining in the anterior cerebellum (Figure 4 E; Figure 4-figure supplement 1 D-E), reminiscent to a previous report that *PV*^*Cre*^ can induce higher levels of transgene expression in anterior than posterior cerebellum (Asrican et al., 2013). Correlating with the expression levels of LZK, there were more Purkinje cells with severe abnormal cell morphology in the anterior cerebellum than the posterior (Figure 4 E). Reduced MAP2 levels were also more noticeable in Purkinje cells located in the anterior than those in the posterior cerebellum (Figure 4 E). Additionally, GFAP staining showed significantly increased astrocyte reactivity in *PV*^*Cre*/+^;*Lzk*^*iOE*/+^ mice from P21 (Figure 4-figure supplement 2 A-B), particularly in areas surrounding Purkinje cells (Figure 4-figure supplement 2 C).

**Figure 4.**
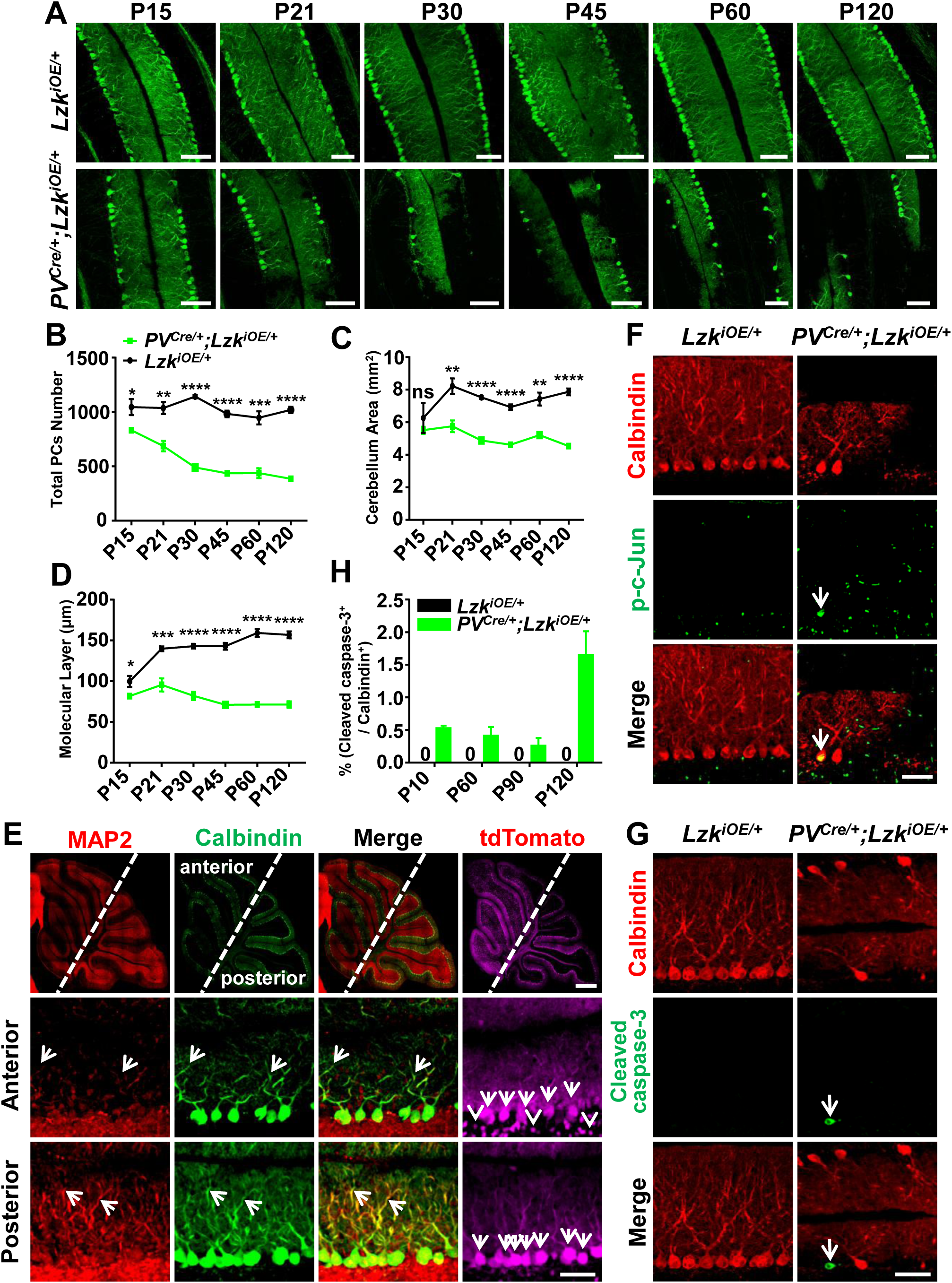
Increasing LZK expression using *PV*^*Cre*^ causes progressive degeneration of Purkinje cells. A. Representative images of Calbindin staining of cerebellar sections of littermates from mating parents *PV*^*Cre*/+^ to *Lzk*^*iOE*/+^ (short for *H11-LSL-Lzk*^*iOE*^ transgene) at indicated postnatal days. Scale bars: 100 μm. B. Quantification of total Purkinje cells in all cerebellar lobules. C. Quantification of cerebellum area, perimeters measured by outlining the outer edge of cerebellum. D. Quantification of the molecular layer thickness in cerebellar lobules V-VI. Color representation for genotypes in C-D is the same as in B. E. Representative images of cerebellar sections of *PV*^*Cre*/+^;*Lzk*^*iOE*/+^ mice at P15, co-immunostained for MAP2 and Calbindin. The dotted lines in top panels mark the boundary of anterior and posterior cerebellum, note higher tdTomato intensity in anterior cerebellum. Images in the bottom two rows show enlarged views of the anterior and posterior cerebellum, with arrows pointing to dendrites of Purkinje cells. Scale bars: 500 μm (upper panel), 100 μm (middle and lower panels). F. Representative images of Purkinje cells of P90 mice co-immunostained for Calbindin and p-c-Jun. Scale bar: 50 μm. G. Representative images of Purkinje cells of P90 mice co-immunostained for Calbindin and cleaved caspase-3. Note that *PV*^*Cre*/+^;*Lzk*^*iOE*/+^ mice have smaller cerebella, hence two rows of Purkinje cells are in view. Scale bar: 50 μm. H. Quantification of the percentage of cleaved caspase-3^+^ Purkinje cells in total Purkinje cells in *Lzk*^*iOE*/+^ and *PV*^*Cre*/+^;*Lzk*^*iOE*/+^ mice from P10 to P120. n = 3 per genotype at each time point. (B-D, H). n ≥ 3 per genotype; data shown are means ± SEM. Statistics for B, C, D, H: Student’s unpaired t-test; ns, no significant; *, p<0.05, **, p<0.01, ***, p<0.001, ****, p<0.0001.

We addressed whether Purkinje cell degeneration caused by LZK involved JNK activation and induction of apoptosis. While no detectable p-c-Jun was observed in Purkinje cells of P90 control mice, LZK activation increased p-c-Jun in Purkinje cells (Figure 4 F). Cleaved caspase-3 immunoreactivity co-localized with Purkinje cells in *PV*^*Cre*/+^;*Lzk*^*iOE*/+^ mice, but not in control mice (Figure 4 G-H). Thus, while elevating LZK expression triggers JNK activation and caspase mediated apoptosis, Purkinje cells undergo a slow degeneration process. These data suggest differential regulation of the signaling network induced by DLK and LZK activation.

### Purkinje cell degeneration induced by LZK overexpression is attenuated by loss of CELF2, a regulator of *Mkk7* alternative splicing

Biochemical studies have shown that two MAP2K, MKK4 and MKK7 act downstream of DLK and LZK to activate JNK (Hirai et al., 2011; Le Pichon et al., 2017; Huang et al., 2017; Ikeda et al., 2001a; Chen et al., 2016b; Ikeda et al., 2001b; Holland et al., 2016; Merritt et al., 1999). However, *in vivo* evidence for how each MAP2K contributes to DLK and LZK induced signal transduction cascade in neurons is limited (Yang et al., 2015). Recent studies of T-cell activation have reported that the activity of MKK7 is regulated through alternative splicing of its exon 2, which encodes a small peptide within the JNK docking site in MKK7 (Martinez et al., 2015) (Figure 5-figure supplement 1 A). During T-cell activation, the RNA splicing factor CELF2 promotes skipping of this exon, favoring the production of a short isoform of MKK7 that has high potency to activate JNK (Martinez et al., 2015; Ajith et al., 2016).

To test if this regulation of *Mkk7* alternative splicing has functional significance in neurons, we generated *PV*^*Cre*/+^;*Celf2*^*fl/fl*^;*Dlk*^*iOE*/+^ and *PV*^*Cre*/+^;*Celf2*^*fl/fl*^;*Lzk*^*iOE*/+^ mice, along with *PV*^*Cre*/+^;*Celf2*^*fl/fl*^ control mice (Figure 5-figure supplement 1 C). By gross animal appearance and morphology of Purkinje cells, *PV*^*Cre*/+^;*Celf2*^*fl/fl*^ mice were indistinguishable from control mice *PV*^*Cre*/+^ or *Celf2*^*fl/fl*^ (Figure 5-figure supplement 1 D-H). Purkinje cell degeneration phenotypes in *PV*^*Cre*/+^;*Celf2*^*fl/fl*^;*Dlk*^*iOE*/+^ remained similar to those in *PV*^*Cre*/+^;*Dlk*^*iOE*/+^ (Figure 5-figure supplement 1 D-F). Deletion of *Celf2* did not alter the levels of DLK (Figure 5-figure supplement 1 I), nor the induction of p-c-Jun (Figure 5-figure supplement 1 J-L). All pups of *PV*^*Cre*/+^;*Celf2*^*fl/fl*^;*Dlk*^*iOE*/+^ had smaller cerebellum (Figure 5-figure supplement 1 G), showed morbidity and low body weight (Figure 5-figure supplement 1 H), and died around P21.

In contrast, *PV*^*Cre*/+^;*Celf2*^*fl/fl*^;*Lzk*^*iOE*/+^ mice showed significantly improved composite behavioral phenotypes, compared to *PV*^*Cre*/+^;*Lzk*^*iOE*/+^ from P30 to P120 (Figure 5 A-E), although the reduced body weight remained in *PV*^*Cre*/+^;*Celf2*^*fl/fl*^;*Lzk*^*iOE*/+^ mice (Figure 5 F). At the cellular level, *Celf2* deletion dramatically reduced Purkinje cell degeneration induced by LZK, with significant improvement in the dendrite morphology at P120 (Figure 5 G-J). *Celf2* deletion also significantly inhibited astrogliosis and microgliosis in P120 *PV*^*Cre*/+^;*Celf2*^*fl/fl*^;*Lzk*^*iOE*/+^ mice, compared to *PV*^*Cre*/+^;*Lzk*^*iOE*/+^ mice (Figure 5-figure supplement 2 A-D). We detected a few cases where microglia appeared to contain dying Purkinje cells in *PV*^*Cre*/+^;*Lzk*^*iOE*/+^ mice but not in *PV*^*Cre*/+^;*Celf2*^*fl/fl*^;*Lzk*^*iOE*/+^ mice (Figure 5-figure supplement 2 C). Consistent with the suppression on Purkinje cell degeneration, *Celf2* deletion also rescued the reduction of MAP2 and NF-200 in the molecular layer of cerebellum caused by LZK activation (Figure 5-figure supplement 3 A-B). Additionally, immunostaining to parvalbumin and neurofilament enabled the visualization of the basket cells, which form specialized structures, the pinceau, onto the axon initial segment (AIS) of Purkinje cells. In *PV*^*Cre*/+^;*Lzk*^*iOE*/+^ mice the pinceau were disorganized, which was suppressed by *Celf2* deletion (Figure 5-figure supplement 3 B-C).

**Figure 5.**
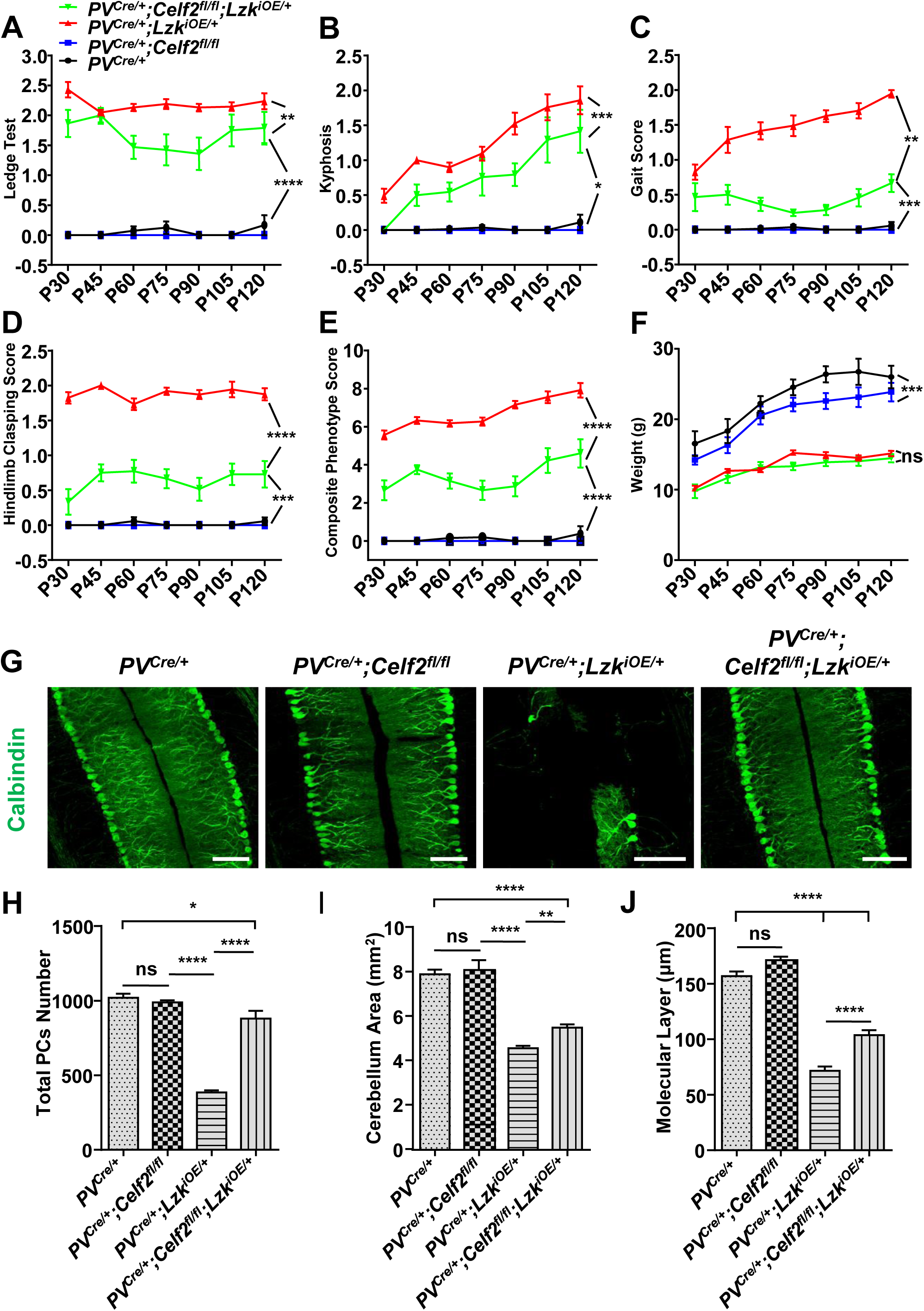
Conditional deletion of the RNA splicing factor *Celf2* using *PV*^*Cre*^ rescues degeneration of Purkinje cells induced by LZK activation. A-E. Quantification of movement phenotypes of mice of indicated genotypes from P30 to P120. *PV*^*Cre*/+^: n ≥ 8; *PV*^*Cre*/+^;*Celf2*^*fl/fl*^: n ≥ 4; *PV*^*Cre*/+^;*Lzk*^*iOE*/+^: n ≥ 7; *PV*^*Cre*/+^;*Celf2*^*fl/fl*^;*Lzk*^*iOE*/+^: n ≥ 5; at each time point. F. Quantification of the body weight of mice of indicated genotypes from P30 to P120. n ≥ 5 per genotype at each time point. Color representation for genotypes in B-F is the same as in A. G. Representative images of Calbindin staining of cerebellar sections from P120 mice of genotypes indicated. Scale bars: 100 μm. H. Quantification of total Purkinje cells in all cerebellar lobules at P120. I. Quantification of cerebellum area at P120. J. Quantification of the molecular layer thickness of P120 mice in cerebellar lobules V-VI. (H-J). n = 3 per genotype; data shown are means ± SEM. Statistics for A-F, H, I, J: One-way ANOVA; ns, no significant; *, p<0.05, **, p<0.01, ***, p<0.001, ****, p<0.0001.

To address whether CELF2 was involved in LZK signaling in Purkinje cells, we examined *Mkk7* exon 2 splicing. By qRT-PCR analysis we detected that *Celf2* deletion reduced the ratio of mRNA of the short isoform (*Mkk7-S*) to the long isoform (*Mkk7-L*) by ∼20% in cerebellum of *PV*^*Cre*/+^;*Celf2*^*fl/fl*^;*Lzk*^*iOE*/+^ mice, compared to *PV*^*Cre*/+^;*Lzk*^*iOE*/+^ mice (Figure 5-figure supplement 1 B). The overall expression levels of LZK in cerebellum were not altered by *Celf2* deletion (Figure 6-figure supplement 1 A-B). We then immunostained for p-c-Jun in cerebellum of P120 mice. LZK overexpression induced p-c-Jun in most of Purkinje cells, and *Celf2* deletion significantly attenuated the intensity of p-c-Jun in Purkinje cells of *PV*^*Cre*/+^;*Celf2*^*fl/fl*^;*Lzk*^*iOE*/+^ mice (Figure 6 A-B). The total number of dying cells marked by TUNEL signals in *PV*^*Cre*/+^;*Celf2*^*fl/fl*^;*Lzk*^*iOE*/+^ mice cerebellum was significantly reduced, compared to that in *PV*^*Cre*/+^;*Lzk*^*iOE*/+^ mice (Figure 6-figure supplement 1 D-E). A few TUNEL signals (approximately one cell / section) were co-localized with tdTomato-labeled Purkinje cells in *PV*^*Cre*/+^;*Lzk*^*iOE*/+^ mice, but not in *PV*^*Cre*/+^;*Celf2*^*fl/fl*^;*Lzk*^*iOE*/+^ mice (Figure 6-figure supplement 1 F). Cleaved caspase-3 signals were rarely detected in Purkinje cells of *PV*^*Cre*/+^;*Celf2*^*fl/fl*^;*Lzk*^*iOE*/+^ mice, compared to *PV*^*Cre*/+^;*Lzk*^*iOE*/+^ mice (Figure 6 C-D). We further assessed expression levels of Bcl-xL (B-cell lymphoma-extra Large), which as a full-length protein prevents caspase activation, but the cleaved product promotes apoptosis (Gross et al., 1999). Western blot analysis of cerebellar protein extracts from P21 mice when minimal Purkinje cell degeneration was detected in *PV*^*Cre*/+^;*Lzk*^*iOE*/+^ showed increased pro-apoptotic cleavage products of Bcl-xL, compared to control samples (Figure 6-figure supplement 1 A, C). All together, these data support a conclusion that *Celf2* deletion attenuates LZK-induced JNK signaling, and provide *in vivo* evidence that MKK7 is a functional mediator of LZK signaling in Purkinje cells.

**Figure 6.**
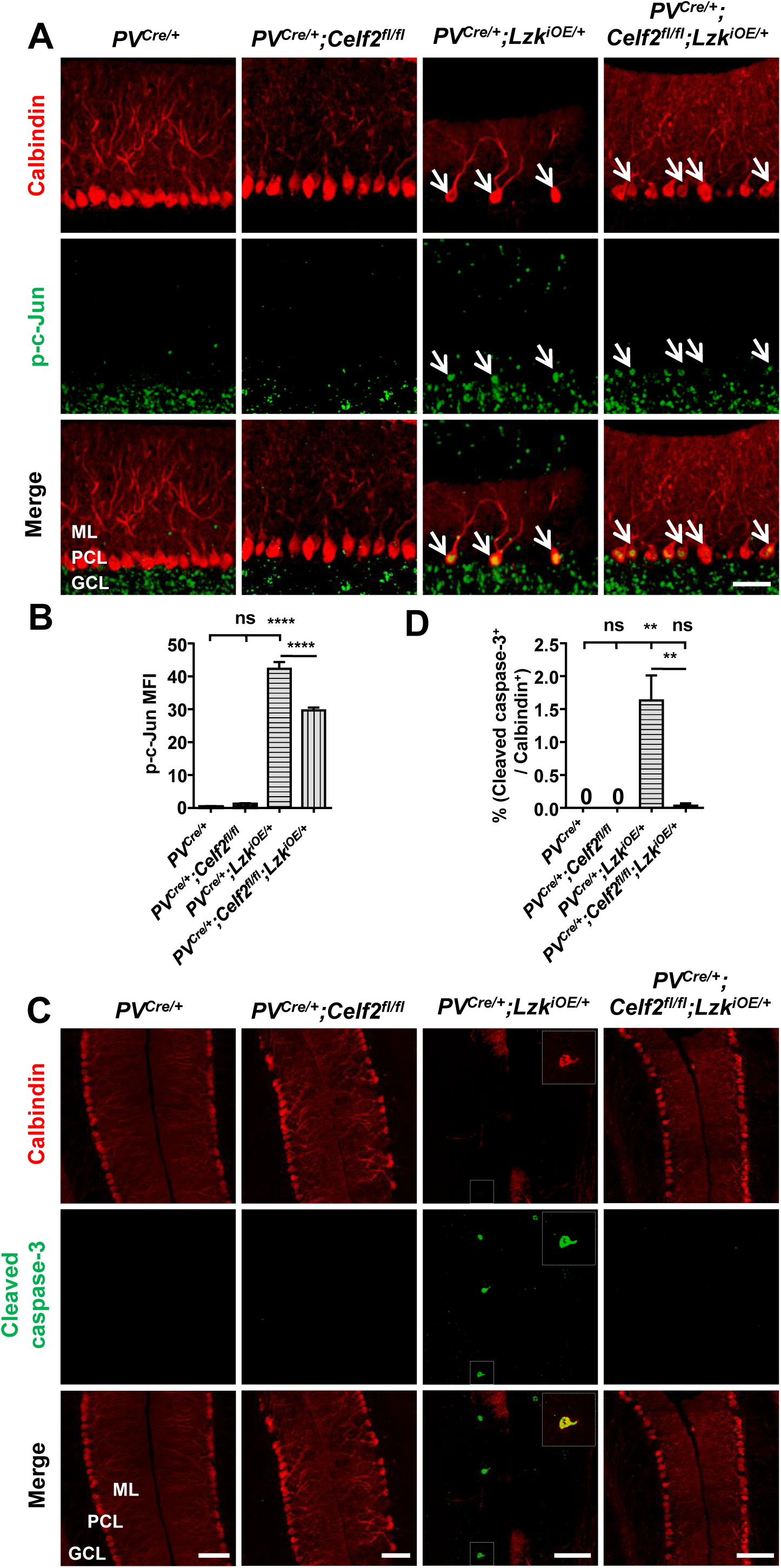
Deletion of *Celf2* in *PV*^*Cre*/+^;*Lzk*^*iOE*/+^ mice reduces levels of p-c-Jun and apoptosis in Purkinje cells. A. Representative images of Purkinje cells, co-immunostained for p-c-Jun and Calbindin, in P120 mice of genotypes indicated. Arrows point to p-c-Jun immunostaining signals in Purkinje cells. ML: Molecular Layer; PCL: Purkinje Cell Layer; GCL: Granule Cell Layer. Scale bar: 100 μm. B. Quantification of the p-c-Jun levels in Purkinje cells of P120 mice. MFI: mean of fluorescence intensity. C. Representative images of Purkinje cells, co-immunostained for cleaved caspase-3 and Calbindin, in P120 mice of genotypes indicated, with enlarged boxes showing cleaved caspase-3^+^ signal in Calbindin labeled Purkinje cells. ML: Molecular Layer; PCL: Purkinje Cell Layer; GCL: Granule Cell Layer. Scale bars: 100 μm. D. Quantification of the percentage of cleaved caspase-3^+^ Purkinje cells in total number of Purkinje cells of P120 mice of genotypes indicated. (B, D). n = 3 per genotype; data shown are means ± SEM. Statistics for B, D: One-way ANOVA; ns, no significant; **, p<0.01, ****, p<0.0001.

### DLK and LZK can induce Purkinje cell degeneration independent of each other

DLK and LZK have a nearly identical kinase domain, and are reported to bind and be co-immunoprecipitated from mouse brain (Pozniak et al., 2013). Recent studies have shown that in injured RGCs or DRGs the two kinases may have redundant or synergistic interactions (Welsbie et al., 2017; Summers et al., 2020). We next addressed whether the Purkinje cell degeneration caused by elevating DLK or LZK activity depends on the presence of one another by analyzing *PV*^*Cre*/+^;*Lzk*^*KO/KO*^;*Dlk*^*iOE*/+^ and *PV*^*Cre*/+^;*Dlk*^*fl/fl*^;*Lzk*^*iOE*/+^ mice.

In *PV*^*Cre*/+^;*Lzk*^*KO/KO*^;*Dlk*^*iOE*/+^ mice, Purkinje cell degeneration and astrogliosis proceeded temporally and spatially similar to that of *PV*^*Cre*/+^;*Dlk*^*iOE*/+^ (Figure 7 A-E). The elevated p-c-Jun levels in Purkinje cells in *PV*^*Cre*/+^;*Dlk*^*iOE*/+^ mice at P10 were not affected by deleting *Lzk* (Figure 7-figure supplement 1 A-B). All pups of *PV*^*Cre*/+^;*Lzk*^*KO/KO*^;*Dlk*^*iOE*/+^ died by P21. Conversely, the *PV*^*Cre*/+^;*Dlk*^*fl/fl*^;*Lzk*^*iOE*/+^ mice resembled *PV*^*Cre*/+^;*Lzk*^*iOE*/+^ mice in the progressive degeneration of Purkinje cells. Both *PV*^*Cre*/+^;*Dlk*^*fl/fl*^;*Lzk*^*iOE*/+^ and *PV*^*Cre*/+^;*Lzk*^*iOE*/+^ mice had low body weight (Figure 7-figure supplement 1 C) and small cerebellum area at P60 (Figure 7 I), compared to control mice. Immunostaining with Calbindin and GFAP antibodies showed that removing *Dlk* did not alter Purkinje cell degeneration or astrogliosis caused by LZK expression (Figure 7 F, H, J). The levels of p-c-Jun induced by LZK expression in Purkinje cells remained comparable, with or without endogenous DLK (Figure 7-figure supplement 1 D-E). Together, these data show that targeted activation of each kinase induces Purkinje cell degeneration largely independent of each other.

**Figure 7.**
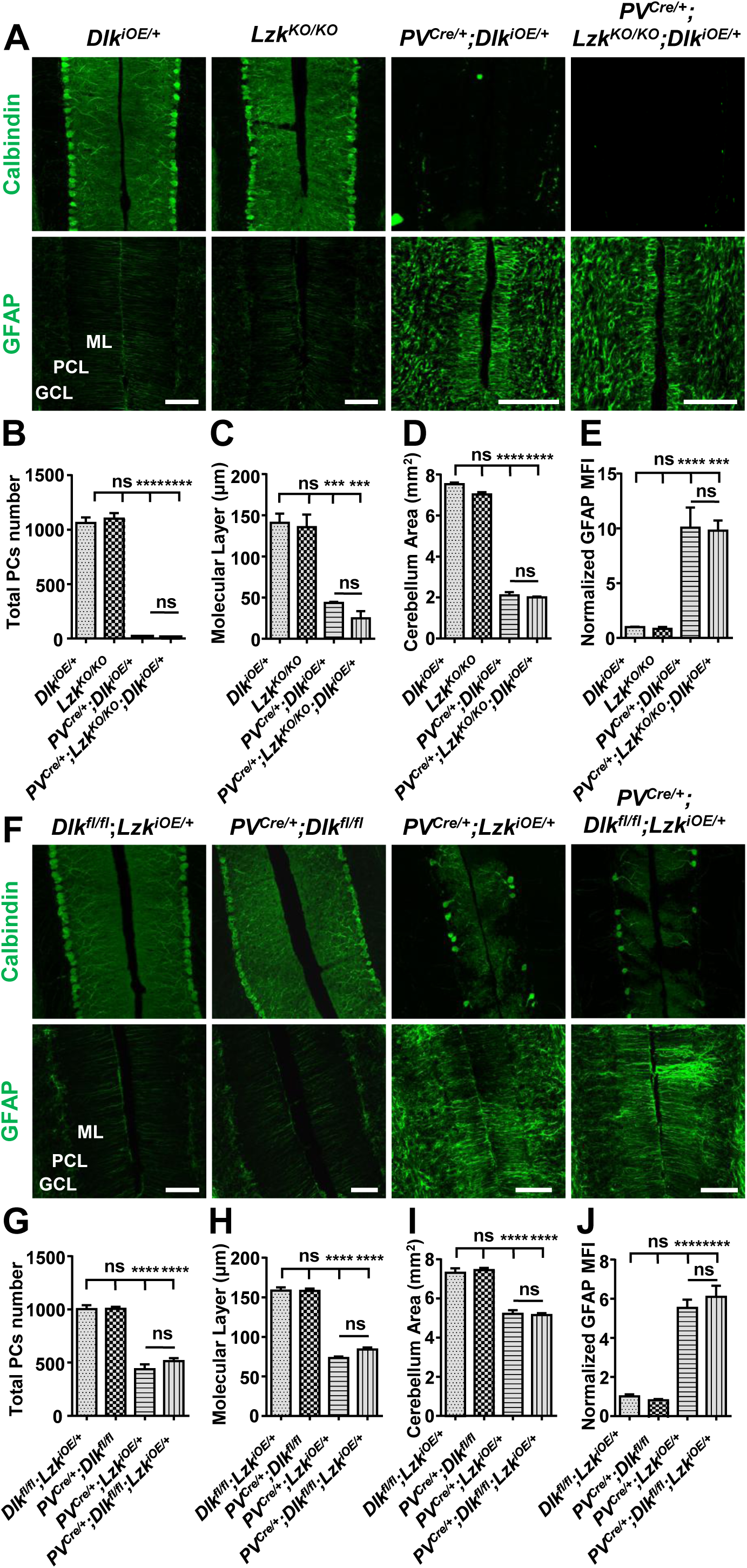
DLK and LZK can induce Purkinje cell degeneration independent of each other. A. Representative images of Purkinje cells of P21 mice of genotypes indicated, immunostained for Calbindin and GFAP. ML: Molecular Layer; PCL: Purkinje Cell Layer; GCL: Granule Cell Layer. Scale bars: 100 μm. B. Quantification of total number of Purkinje cells in all cerebellar lobules at P21. C. Quantification of the molecular layer thickness in cerebellar lobules V-VI of P21 mice. D. Quantification of the cerebellum area of P21 mice. E. Quantification of normalized GFAP levels in cerebellum of P21 mice. MFI: mean of fluorescence intensity. F. Representative images of of Purkinje cells of P60 mice of genotypes indicated, immunostained for Calbindin and GFAP. ML: Molecular Layer; PCL: Purkinje Cell Layer; GCL: Granule Cell Layer. Scale bars: 100 μm. G. Quantification of total number of Purkinje cells in all cerebellar lobules of P60 mice. H. Quantification of the molecular layer thickness in cerebellar lobules V-VI of P60 mice. I. Quantification of the cerebellum area of P60 mice. J. Quantification of normalized GFAP levels in cerebellum of P60 mice. MFI: mean of fluorescence intensity. (B-E, G-J). n = 3 per genotype; data shown are means ± SEM. Statistics for B-E, G-J: One-way ANOVA; ns, no significant; ***, p<0.001, ****, p<0.0001.

## Discussion

In this study, we have used cerebellar Purkinje cells to gain a systematic understanding of the function of DLK and LZK, two closely related kinases that have emerged as key players in neural protection under injury and disease (Adib et al., 2018; Jin and Zheng, 2019; Farley and Watkins, 2018). We employed both conditional KO and transgenic mouse models to manipulate levels of DLK and LZK expression. We find that while deleting one or both kinases in Purkinje cells postnatally did not affect neuronal development and animal health, activating DLK or LZK, through elevating their expression, causes Purkinje cell degeneration. Our Cre-inducible DLK and LZK transgenes have the same design and are inserted in the same *H11* locus to avoid position effect on transgene expression. Despite the similarly targeted transgenes, we found that DLK elevation triggers rapid degeneration of Purkinje cells, while LZK elevation causes slow degeneration. Each kinase activates JNK signaling, measured by increased phosphorylated c-Jun, and induces apoptosis. Each kinase can induce neuron degeneration in the absence of the other. Importantly, we show that deletion of *Celf2* strongly attenuates Purkinje cell degeneration caused by LZK, but not DLK, activation, providing further evidence for a signaling pathway-specific effect for each kinase activation rather than a generic, secondary effect of overexpressing any kinase. As Purkinje cells and cerebellum are not essential for animal viability, we interpret that the lethality of *PV*^*Cre*/+^;*Dlk*^*iOE*/+^ pups is likely due to disruption of other parvalbumin-expression neurons, with the underlying basis remaining to be addressed in future studies. All together, these data demonstrate the utility of our transgenic mice for dissecting cell-type specific roles of these kinases and their signaling pathways.

DLK and LZK share a kinase domain that is ∼90% identical and can activate the JNK signaling pathway through two MAPKK, MKK4 or MKK7 (Hirai et al., 2011; Le Pichon et al., 2017; Huang et al., 2017; Ikeda et al., 2001a; Chen et al., 2016b; Ikeda et al., 2001b; Holland et al., 2016; Merritt et al., 1999). Several studies have supported MKK4 as a major mediator for DLK in RGCs and DRGs (Yang et al., 2015). Currently, little is known which MAPKK mediates LZK signaling. Our data show that DLK activation in Purkinje cells leads to robust JNK signaling, compared to LZK activation. The observation that *Celf2* deletion did not affect any phenotypes caused by DLK activation could be due to a combination of the strong JNK activation and the rapid time course of cell death. In contrast, deletion of CELF2 significantly reduced the activation of c-Jun and almost completely rescued the Purkinje cell degeneration caused by LZK activation. These data are consistent with the role of *Celf2* in regulating alternative splicing of *Mkk7* (Martinez et al., 2015), and support that MKK7 is a functional downstream kinase for LZK *in vivo*.

Numerous studies have revealed roles of DLK in axon growth, regeneration and/or degeneration (Tedeschi and Bradke, 2013; Jin and Zheng, 2019). However, not much is known about roles of LZK in neurons. Our data show that LZK activation decreased neurofilament levels in the molecular layer of cerebellum and caused disorganization of the pinceau at the axon initial segment of Purkinje cell. DLK is known to regulate microtubule stability (Simard-Bisson et al., 2017; Valakh et al., 2015; Hirai et al., 2011), and several microtubule-associated proteins such as SCG10, DCX and MAP2 are JNK substrates (Chang et al., 2003; Gdalyahu et al., 2004; Tararuk et al., 2006; Björkblom et al., 2005). We find that both DLK and LZK overexpression significantly decreased MAP2 levels in dendrites of Purkinje cells, and that *Celf*2 deletion restored MAP2 levels in *PV*^*Cre*/+^;*Celf2*^*fl/fl*^;*Lzk*^*iOE*/+^ mice. In addition, DLK activation caused dendrite swelling of Purkinje cells, and many of the swollen dendrites also had cleaved caspase-3 signals. Activated caspase-3 in dendrites has been shown to cause cleavage of microtubules and local pruning of dendrites and spines (Ertürk et al., 2014; Khatri et al., 2018). These data are consistent with known roles of JNK regulation of microtubule associated proteins.

Taken together, our findings indicate that DLK-induced signal transduction cascade triggers a strong response under injury or other stress, while LZK induces modest activation of JNK and apoptosis, which may manifest in chronic neurodegenerative diseases. Besides the kinase domain and leucine-zipper domain, both MAP3Ks have large uncharacterized C-terminus, which may play significant roles in regulating the signaling strength of each protein. There is mounting evidence in the literature on the upregulation of the DLK and LZK signaling pathway in CNS injury and neurodegeneration (Watkins et al., 2013; Welsbie et al., 2013; Joy et al., 2019; Chen et al., 2008; Le Pichon et al., 2017; Huang et al., 2017; Chen et al., 2018), indicating that altered signaling of this pathway may be a prevalent feature in CNS injury and diseases. Along this line, genetic studies of DLK in both invertebrate and vertebrate species revealed prominent developmental defects with DLK activation rather than inactivation (Zhen et al., 2000; Schaefer et al., 2000; Wan et al., 2000; Grill et al., 2016). As DLK and LZK activity exhibits high cell-type and context-dependent specificity, our transgenic mice offer valuable gain of function models to study their signaling pathways with the ease for temporal and spatial manipulation. The knowledge learned will advance our understanding of how diverse neuronal types respond to insults to the nervous system.

## Materials and methods

### Mice

All animal protocols were approved by the Animal Care and Use Committee of the University of California San Diego. Wild-type C57BL/6J mice and *PV*^*Cre*^ mice (Stock No: 017320) were purchased from The Jackson Laboratory.

*Lzk* knockout mice were generated in the UCSD Transgenic and Knockout Mouse Core, using CRISPR-Cas9 technology (Ran et al., 2013). Briefly, sgRNA sequences targeted to the kinase domain were designed using online tools (http://crispr.mit.edu) (Table 1). The selected sgRNAs were annealed, and then cloned into PX330 backbone digested with BbsI. Effectiveness of sgRNAs was tested using Surveyor nuclease assay (Surveyor Mutation Detection Kit, IDT, 706020). To make sgRNAs, DNA fragments containing T7 promoter followed by sgRNA were first amplified using primers YJ12532-12535. The purified DNAs were then in vitro transcribed using MEGAscript T7 Transcription Kit (Invitrogen, AMB13345), and the resulting transcripts were purified using MEGAclear-96 Transcription Clean-Up Kit (Invitrogen, AM1909). The sgRNAs and Cas9 mRNA were injected into zygotes from C57BL6, which were then implanted into the CD1 surrogate mothers. Two KO mouse lines were obtained and the one containing a deletion of the entire kinase domain was used in this study. *Lzk*^*fl*^ and *Dlk*^*fl*^ mice were reported in (Chen et al., 2016b). *Dlk*^*fl*^ mice were a kind gift of Dr. Lawrence B. Holzman (Univ. Penn). *Celf2*^*fl*^ mice were described previously (Chen et al., 2016a). Primers for genotypes are listed in Table 2.

**Table 1:**
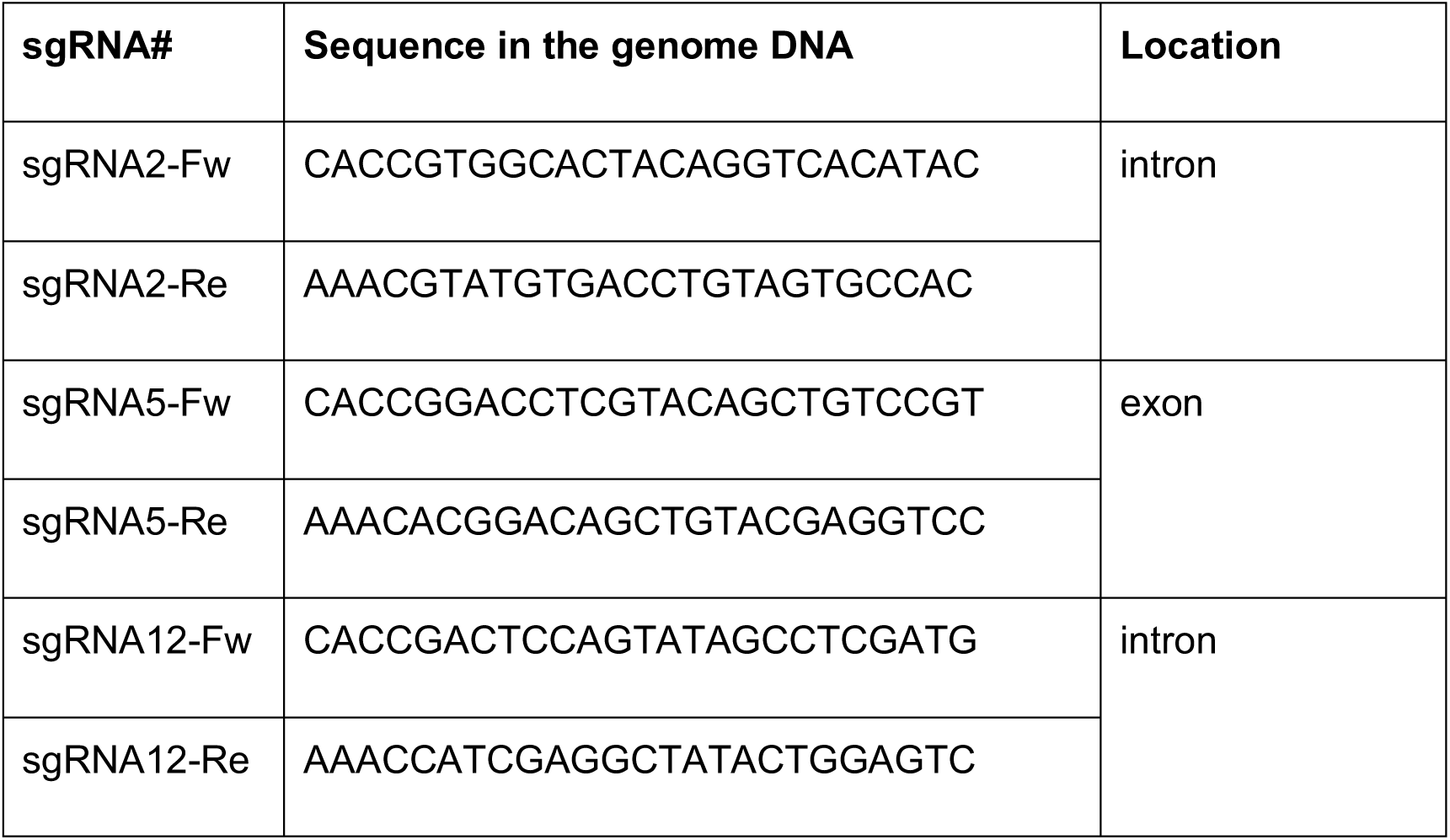
sgRNA sequences used to generate LZK CRISPR KO mice.

**Table 2:**
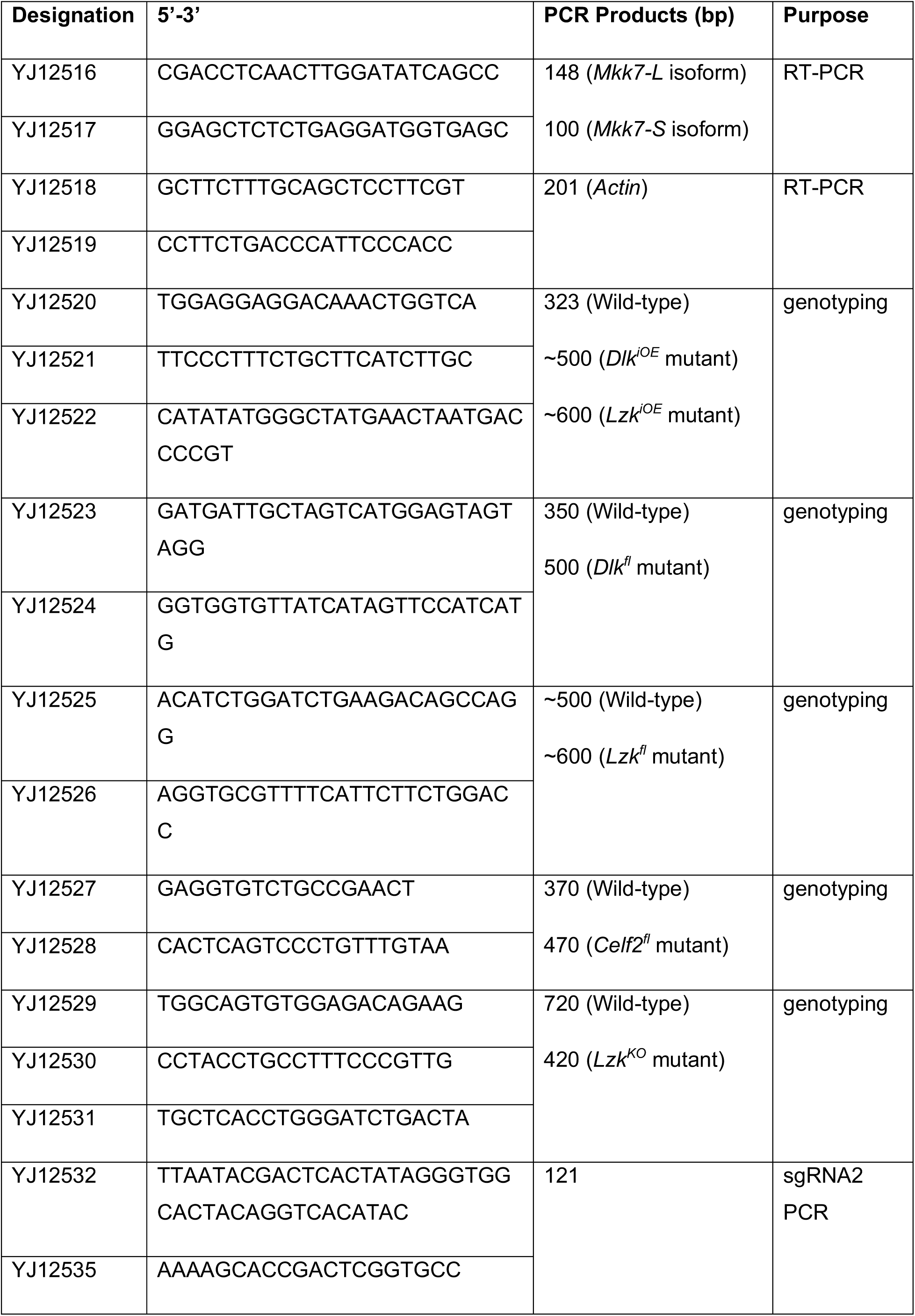
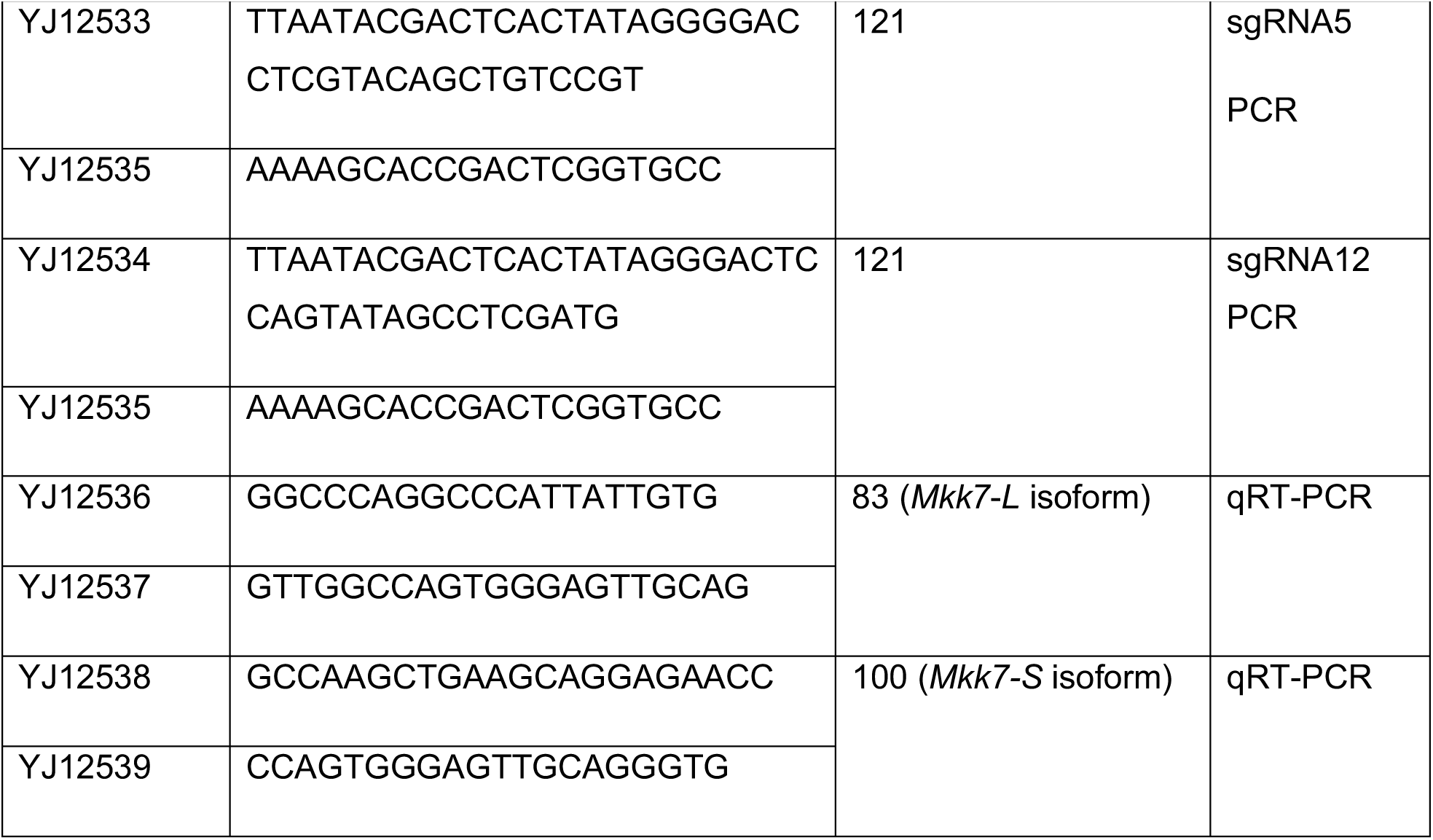
Primers used in this study.

Transgenic conditional overexpressing *Dlk*^*iOE*^ and *Lzk*^*iOE*^ mice were made by Applied StemCell, Inc (Milpitas, CA), using TARGATT™ Technology (Tasic et al., 2011). A mixture of plasmid pBT378-LSL-3X Flag-*Dlk*-T2A-tdTomato, or pBT378-LSL-1X Flag-*Lzk*-T2A-tdTomato DNA, and in vitro transcribed ϕC31 integrase mRNA was microinjected into the pronucleus of zygotes from a FVB strain that has the Att recombination landing site inserted in *H11* locus, which were then implanted into the CD1 surrogate mothers. The founder heterozygous mice were bred three times to pure C57BL/6J background.

### Histology and immunocytochemistry

Mice were transcardially perfused with 0.9% saline solution and then 4% paraformaldehyde (PFA) in phosphate-buffered saline (PBS) (pH 7.2-7.4). Brains were dissected, post-fixed in 4% PFA overnight at 4°C, and then transferred to 30% sucrose in PBS, prior to embedding using O.C.T compound (Fisher Healthcare, 4585) on dry ice. 25 μm thick sagittal sections were collected on a cryostat (Leica, CM1850) into PBS with 0.01% NaN_3_. For histology analysis, free floating tissue sections were loaded to the slide, and then sequentially stained with hematoxylin and eosin (H&E staining Kit, abcam, ab245880). For immunostaining, free floating tissue sections were washed twice in PBS with 0.2% Triton X-100, blocked with 5% goat serum in PBS with 0.4% Triton X-100 for 2 hours at room temperature, then incubated with primary antibodies (Table 3) diluted in PBS with 0.2% Triton X-100 and 2% goat serum overnight at 4°C. Alexa Fluor 488-conjugated and Alexa Fluor 647-conjugated secondary antibodies (Invitrogen) were incubated for 2 hours at room temperature, followed by staining with DAPI (14.3 μM in PBS, Thermo Fisher Scientific, D1306) for 10 min. Stained sections were mounted with prolong diamond antifade mountant (Thermo Fisher Scientific, P36970).

**Table 3:**
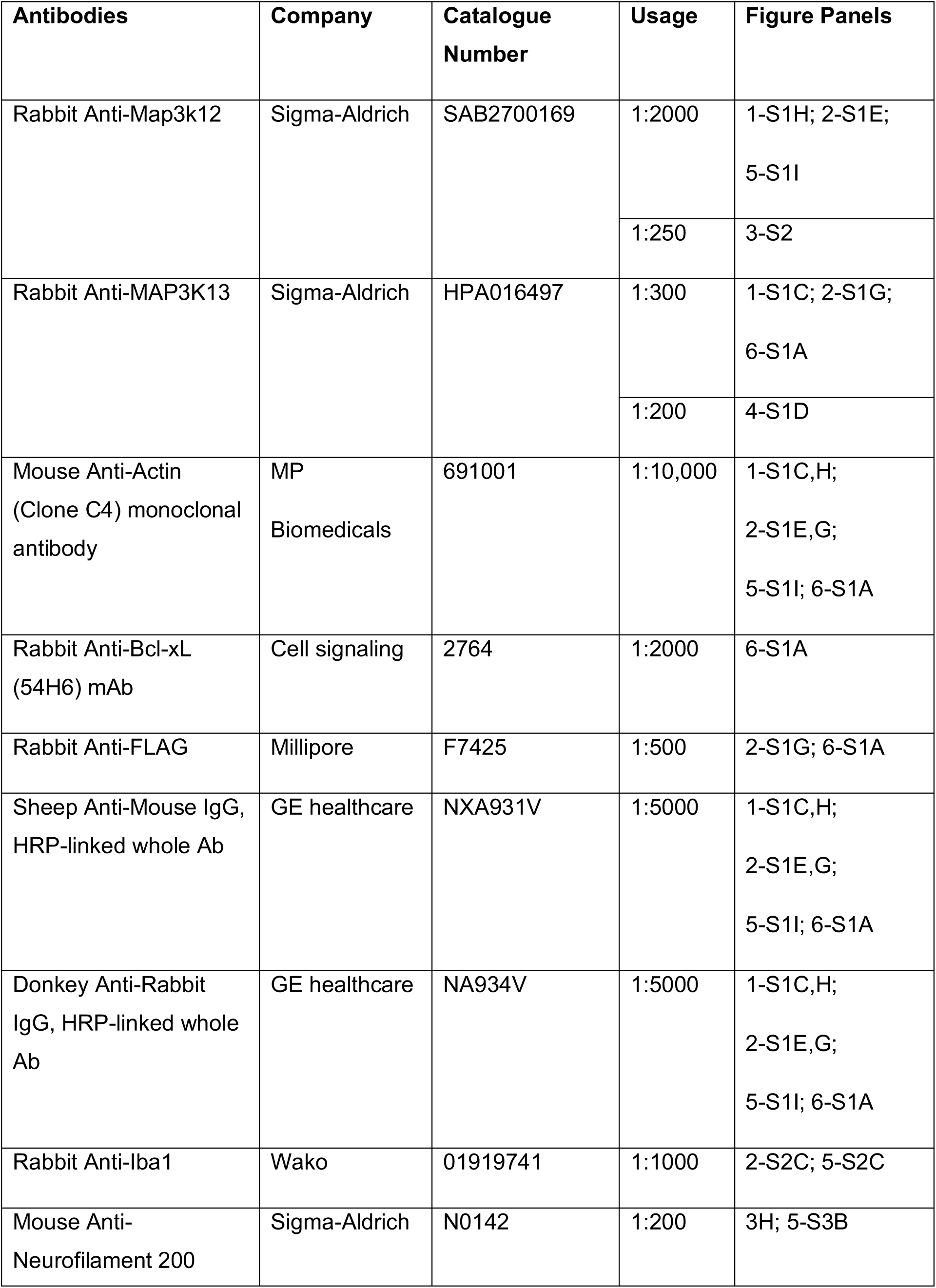
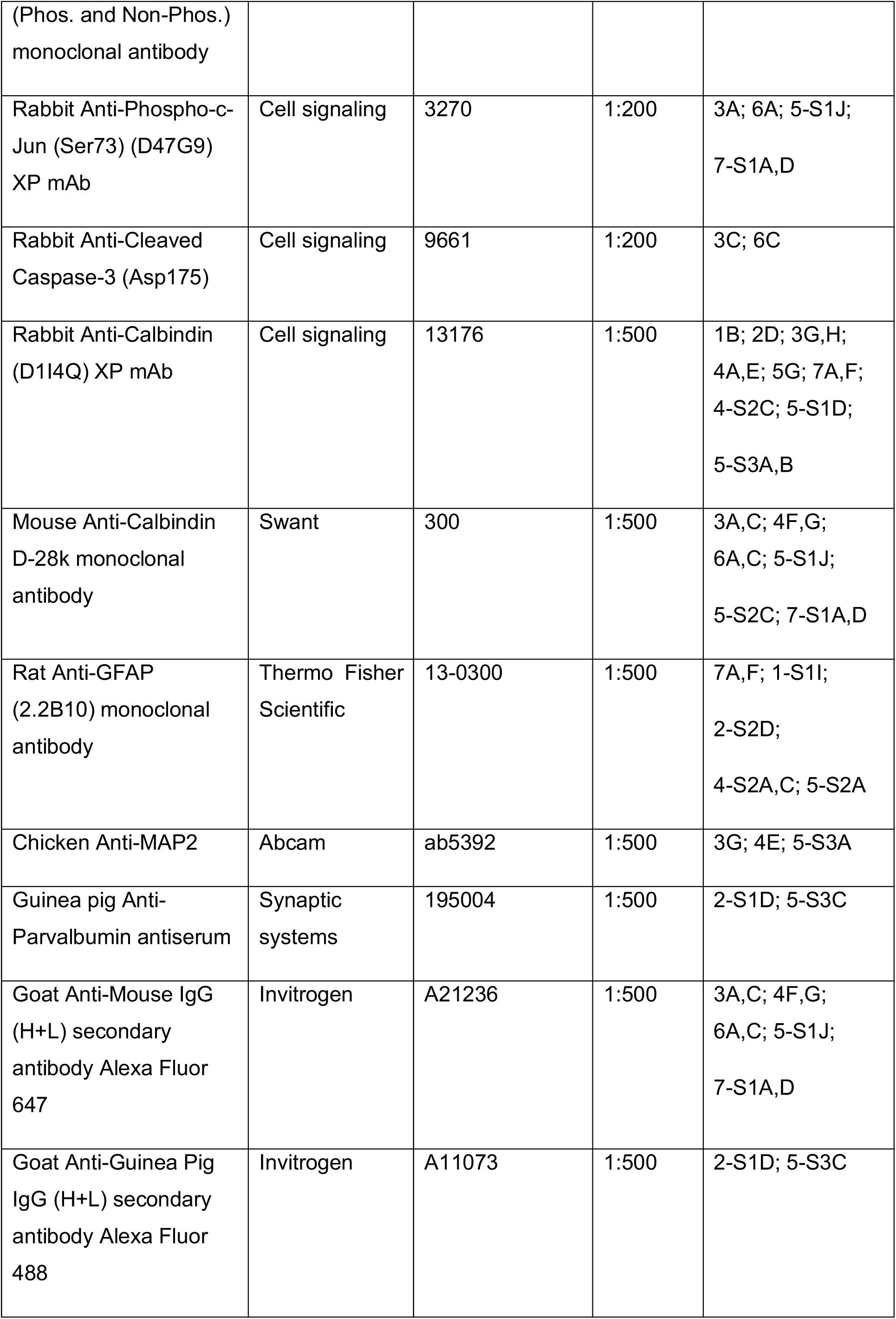
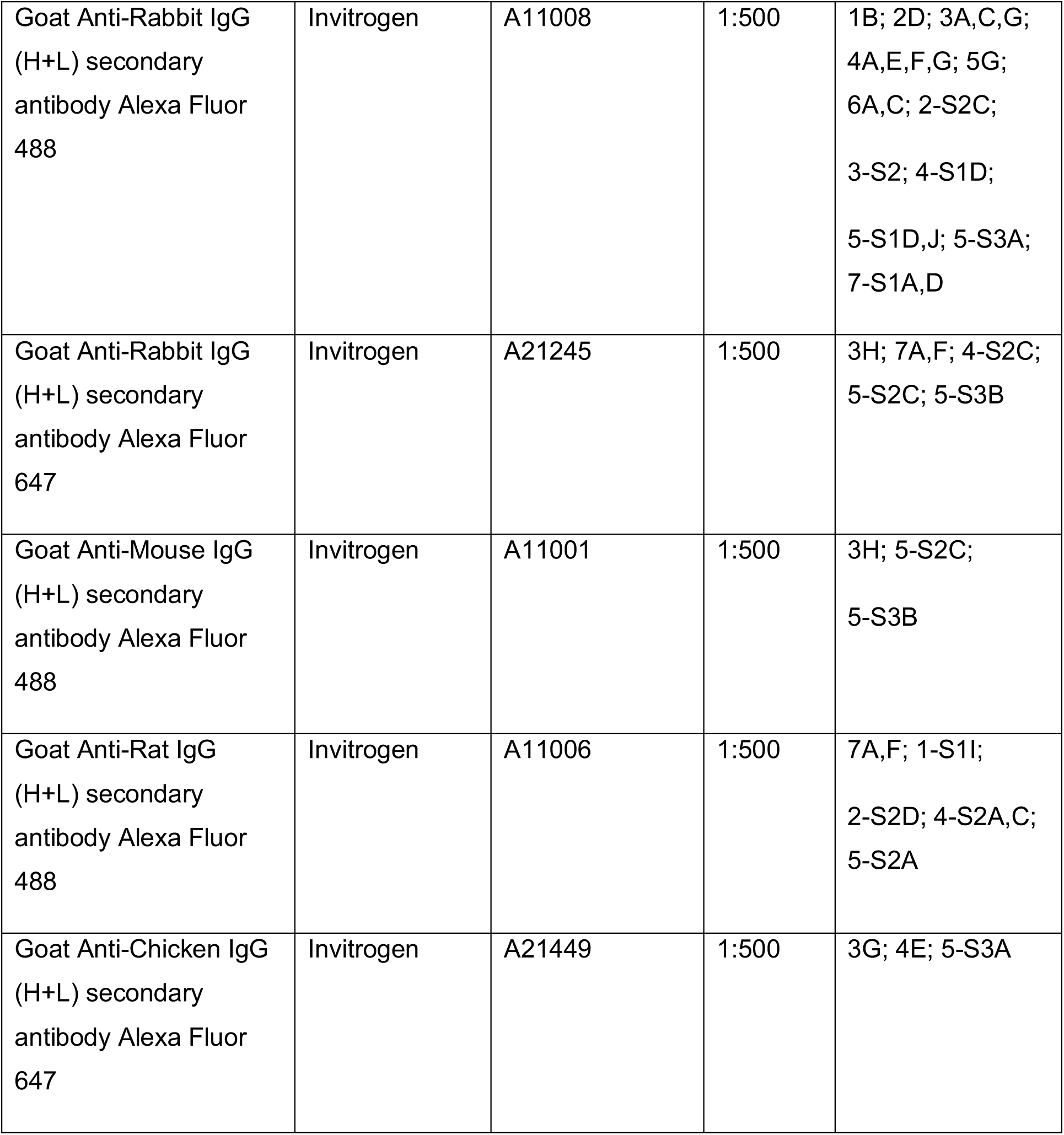
Antibodies used in this study.

### TUNEL staining

The DeadEnd Fluorometric TUNEL System (Promega, G3250) was used with a modified protocol. Free floating tissue sections were washed twice in PBS, and then loaded to the slide. The slides were dried at 65°C for 5 min, then immersed into PBS with 0.5% Triton X-100 and incubated at 85°C for 20 min, followed by three times rinsing with PBS. The slides were incubated with equilibration buffer at room temperature for 5 min, and then incubated with reaction mix (equilibration buffer: nucleotide mix: rTdT enzyme = 45:5:1) at 37°C for 1 hour in a humidity chamber. The reactions were stopped by incubating the slide with 2X saline-sodium citrate (SSC) buffer at room temperature for 10 min. After three washes with PBS, the slides were incubated with DAPI (14.3 μM in PBS) at room temperature for 15 min, followed by three washes with ddH_2_O, and then mounted with prolong diamond antifade mountant (Thermo Fisher Scientific, P36970).

### Image acquisition and analysis

The slides of H&E staining were scanned with Nanozoomer 2.0-HT digital slide scanner (Hamamatsu) in brightfield at 20X magnification. The images were processed using NDP.view2 Viewing software (Hamamatsu) and ImageJ software (NIH). Fluorescence images of paired WT and mutant samples were acquired on a Zeiss LSM 710 confocal microscope. The images were taken as Z-stack under identical settings, and the maximum intensity projection images were processed using ImageJ software (NIH). For image quantification, three midline parasagittal sections per brain and at least 3 brains per genotype of given age were analyzed and data was averaged. Cells were counted using the cell counter plugin for ImageJ (NIH). Analyses of cell numbers for Calbindin^+^ PCs, p-c-Jun^+^ PCs, dendrite swelling^+^ PCs or cleaved caspase-3^+^ PCs were performed by counting the soma of each PC in the entire lobules. The thickness of the molecular layer visualized by Calbindin staining was assessed for lobule V/VI in midline sections by measuring the perpendicular distance from the molecular layer-facing edge of a Purkinje cell soma to the outer edge of the molecular layer. Cerebellum area was calculated by outlining the perimeter of the outer edges of the sagittal sections of cerebellum. TUNEL^+^ cells were counted by analyzing particles after adjustment of threshold and watershed. The particles with area larger than 8 μm^2^ were measured. TUNEL^+^ cell density was calculated the number of TUNEL^+^ cells in entire cerebellum divided by entire cerebellum area. GFAP or IBA1 immunofluorescence intensity density was calculated by dividing the GFAP or IBA1 immunofluorescence intensity of entire cerebellum by the entire cerebellum area. For p-c-Jun immunofluorescence intensity quantification, 50 sampling area surrounding a single p-c-Jun^+^ nucleus (ROI being 347.543 μm^2^) per section were measured. For LZK immunofluorescence intensity quantification, 30 sampling area surrounding a single soma of Purkinje cells (ROI being 352.943 μm^2^) were measured in the region of interest per section. Integrated density was averaged after subtraction of background signal and adjustment of threshold.

### Immunoprecipitation and western blotting

Dissected cerebella from mice of indicated age were homogenized in an appropriate volume of cell lysis buffer (50 mM Tris.Cl (pH 7.4), 1% Triton X-100, 0.1% SDS, 1 mM EDTA (pH 7.0), 150 mM NaCl, 1% n-Octyl β-D-glucopyranoside, 1 x protease inhibitor cocktail (Roche, 05892970001)) using TissueRuptor II (QIAGEN, 9002756), then lysed for 1 hour on ice, and cleared by centrifugation at 13,000 rpm for 10 min at 4°C. Supernatants were collected and protein concentrations were determined by Pierce BCA protein assay kit (Thermo Fisher Scientific, 23225). For immunoprecipitation experiments, antibody-bound beads were prepared using 2 μg rabbit anti-MAP3K13 polyclonal antibody (Sigma-Aldrich, HPA016497) in 800 μl of lysis buffer with 50 μl of 50% Protein G agarose bead slurry, with gentle rotation at 4°C for 1 hour. ∼1 mg protein lysates were pre-cleared with 50 μl of 50% Protein G agarose bead slurry, then incubated with the antibody-bound beads overnight at 4°C. The beads were washed three times with lysis buffer, and then resuspended in 60 μl lysis buffer and 20 μl 4 x Laemmli Sample Buffer (Bio-RAD, 161-0747), heat shocked in a thermomixer (Eppendorf) at 95°C for 10 min and analyzed by western blotting. Immunoprecipitated samples were separated by SDS-PAGE using Any kD Mini-PROTEAN TGX Precast Protein Gels (Bio-Rad, 4569034), and then blotted to a PVDF membrane (0.2 μm, Bio-RAD, 1620177) by Mini Trans-Blot Cell (Bio-RAD, 170-3930) at 100 mA for 1 hour. Blots were blocked in 10% non-fat dry milk in PBST (PBS with 0.05% Tween-20) for 1 hour at room temperature, and then incubated with an appropriate concentration of primary antibody in 1% non-fat BSA in PBST at 4°C for overnight. Afterwards, the membrane was incubated with Horseradish Peroxidase (HRP)-conjugated secondary antibody (GE healthcare, NXA931V or NA934V) in 1% non-fat BSA in PBST at room temperature for 1 hour, followed by detection using enhanced chemiluminescence (ECL) reagents (GE Healthcare, RPN2106).

### RNA extraction, RT-PCR and qRT-PCR

Total RNA from mouse cerebellum was extracted using TRIzol (Invitrogen, 15596018) following the manufacturers’ protocols. First strand cDNA was reverse-transcribed using SuperScript IV (Invitrogen, 18091050). qPCR was run on Bio-Rad CFX96 Touch Real-Time PCR Detection System with iQ SYBR Supermix (Bio-Rad, 170-8882). Data were analyzed using CFX manager (Bio-Rad).

### Animal behavior tests

Mouse behavioral analysis was scored in a genotype blind manner following the protocol described in (Guyenet et al., 2010). Briefly, ledge walking, hind limb clasping, gait, and kyphosis were scored with a scale of 0-3 in each category, resulting in total score of 0-12 points for all four measures at P30, P45, P60, P75, P90, P105 and P120. A score of 0 represents absence of the relevant phenotype and 3 represents the most severe phenotype. Each test was performed 3 times to ensure reproducibility. For data analysis, the score was calculated for each measure by taking the mean of the three measurements in each mouse.

### Statistics

GraphPad Prism 6.0 (GraphPad Software, Inc) was used for all statistical analysis. After assessing for normal distribution, statistical analyses between two groups were calculated with the two-tailed t-test for normally distributed data. For comparison of more than two groups, normally distributed data was calculated with a one-way ANOVA. Asterisks indicate significance with (*) P<0.05, (**) P<0.01, (***) P<0.001, (****) P<0.0001 for all data sets. Graphs show mean values ± standard error of the mean (SEM).

## Acknowledgements

We are grateful to our lab members for valuable discussions. We thank Drs. A. D. Chisholm, S. L. Ackerman and G. Thomas for comments on the manuscript. We thank Dr. L. Holzman (U. Penn) for providing *Dlk*^*fl*^ mice, UCSD Transgenic and Knockout Mouse Core for generating *Lzk*^*KO*^ mice, and UCSD Neuroscience Microscopy Shared Facility (NS047101) for providing imaging support. We thank A. Moore, E. Xu, D. Arakelyan and R. Zarei for technical assistance. This work was partly supported by funds from Howard Hughes Medical Institute, the Craig H. Neilsen Foundation, the Junior Seau Foundation, and the UCSD Kavli Institute of Brain and Mind.

## Author contributions

Y.L., Y.J. conceived of the study and wrote the manuscript. Y.L., E.R., C.S., C.Q., L.Z. performed the experiments and data analyses. All authors contributed to writing, reviewing, editing of the manuscript.

## DECLARIATION OF INTERESTS

The authors declare no competing interests.

## Supplementary figure legends

**Figure 1-figure supplement 1.**
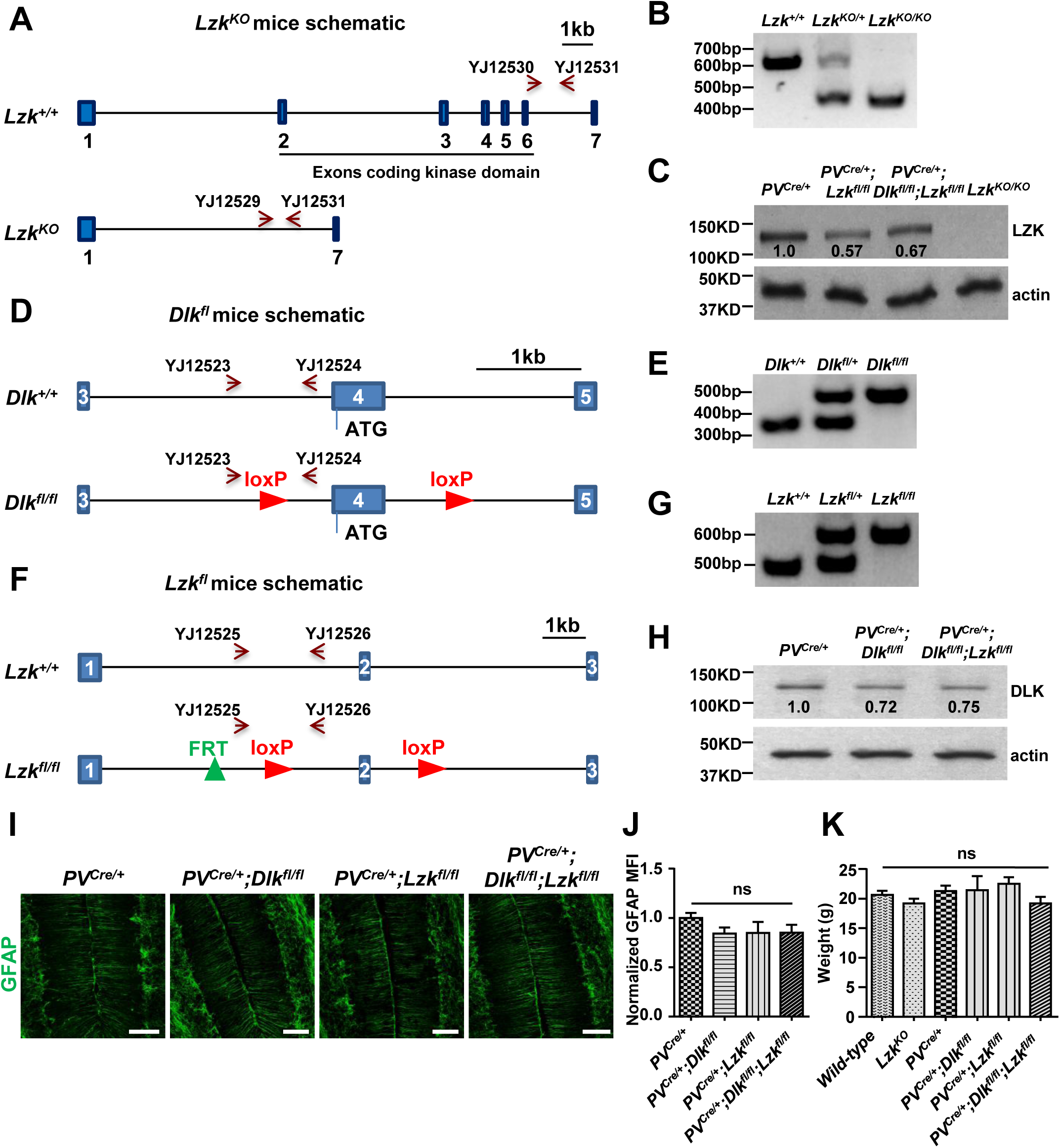
Knockout mice lines and associated evidence. A. *Lzk* KO mice were generated by CRISPR-Cas9 technology using 3 sgRNAs targeted to the kinase domain of LZK. Scale bar: 1kb. B. The homozygotes or heterozygotes of *Lzk*^*KO*^ were identified by PCR using primers YJ12529-12531. C. The LZK protein levels in cerebellum of P60 mice of genotypes indicated were determined by immunoprecipitation and western blot. D. *Dlk*^*fl/fl*^ mice have loxP sites flanking the exon 4 which contains the initiation ATG codon, and Cre-mediated excision results in a null allele. Scale bar: 1kb. E. The homozygotes or heterozygotes of *Dlk*^*fl*^ were identified by PCR using primers YJ12523 and YJ12524. F. *Lzk*^*fl/fl*^ mice have loxP sites flanking exon 2, and Cre-mediated excision results in frameshift, hence a null allele. Scale bar: 1kb. G. The homozygotes or heterozygotes of *Lzk*^*fl*^ were identified by PCR using primers YJ12525 and YJ12526. H. The DLK protein levels in cerebellum of P60 mice of genotypes indicated were determined by western blot. I. Representative images of GFAP staining of cerebellar astrocytes from P60 mice of genotypes indicated. Scale bars: 100 μm. J. Quantification of normalized GFAP levels in cerebellum of P60 mice. n = 3 per genotype. MFI: mean of fluorescence intensity. K. Quantification of the body weight of mice of genotypes indicated at P60. n ≥ 3 per genotype. (J, K). Data shown are means ± SEM. Statistics for J, K: One-way ANOVA; ns. no significant.

**Figure 2-figure supplement 1.**
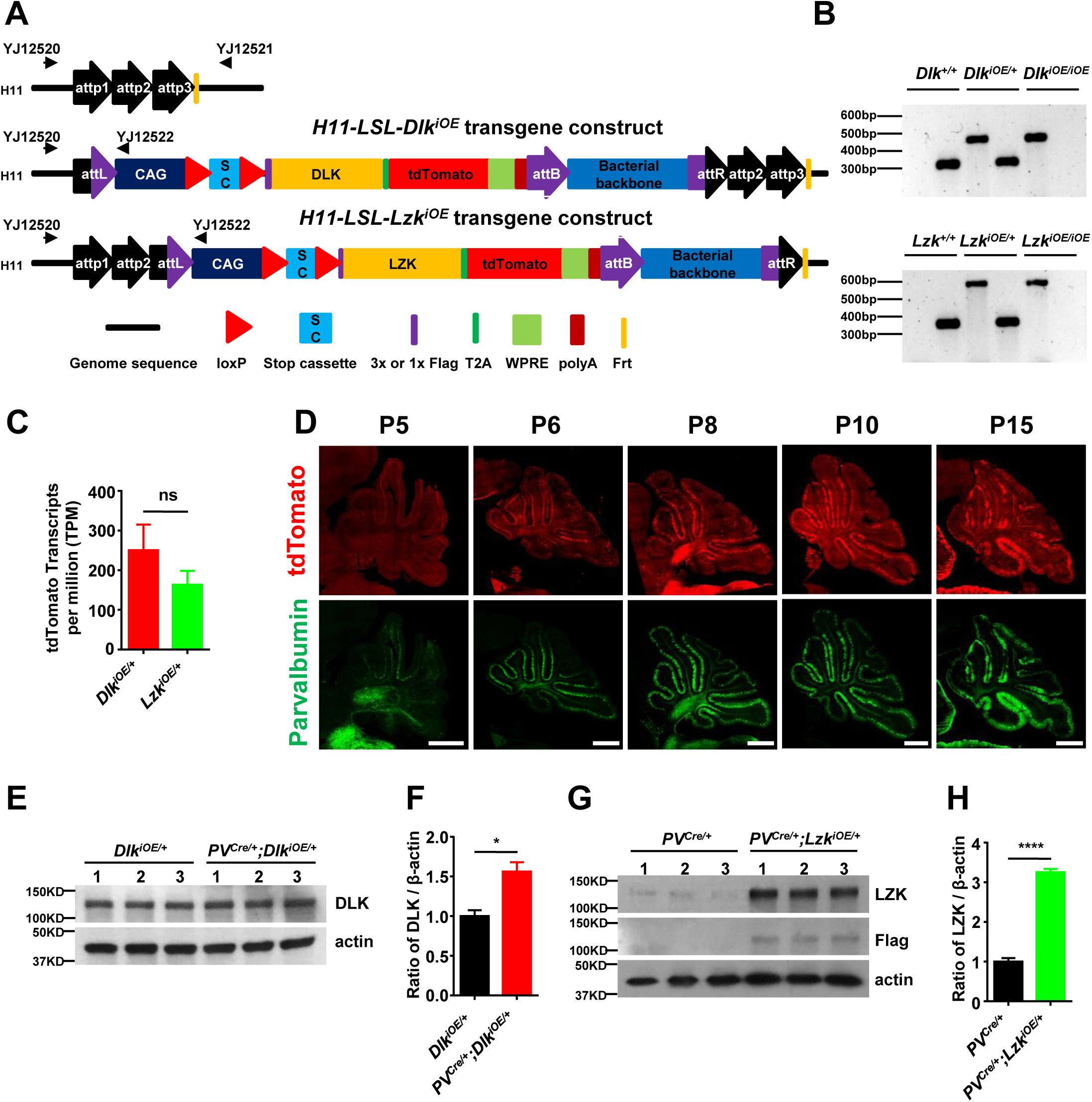
Cre-dependent DLK and LZK expression transgenic mice and associated evidence. A. Illustration of Cre-dependent expression of *Dlk*^*iOE*^ and *Lzk*^*iOE*^ transgenic mice, generated following the protocol described previously (Tasic et al., 2011). 3 x Flag-*Dlk* or 1 x Flag-*Lzk* and tdTomato will be expressed after removal of the stop cassette by Cre recombinase. B. The homozygotes or heterozygotes of *Dlk*^*iOE*^ and *Lzk*^*iOE*^ were identified by PCR using primers YJ12520-12522. C. Graph shows mRNA levels of tdTomato obtained from RNA-sequencing of cerebellum from P15 mice of *Vglut1*^*Cre*/+^;*Dlk*^*iOE*/+^ and *Vglut1*^*Cre*/+^;*Lzk*^*iOE*/+^. Transcript per million (TPM) values represent tdTomato transcript counts normalized by gene length and sequencing depth. Differences in the expression levels of tdTomato between *Vglut1*^*Cre*/+^;*Dlk*^*iOE*/+^ and *Vglut1*^*Cre*/+^;*Lzk*^*iOE*/+^ is not statistically significant based on differential expression analysis using DESeq2 galaxy version 2.11.40.2 (Love et al., 2014). D. Representative images of cerebellar sections of *PV*^*Cre*/+^;*Dlk*^*iOE*/+^ mice immunostained for parvalbumin at indicated postnatal days, along with tdTomato expression from *Dlk*^*iOE*^, showing the effect of Cre recombinase. Scale bars: 500 μm. E. The DLK protein levels in cerebellar extracts from P10 mice of genotypes indicated were determined by western blot. F. Quantification of the ratio of DLK relative to β-actin protein levels. G. The LZK protein levels in cerebellar extracts from P21 mice of genotypes indicated were determined by immunoprecipitation and western blot. H. Quantification of the ratio of LZK relative to β-actin protein levels. (C, F, H). n = 3 per genotype; data shown are means ± SEM. Statistics for C, F, H: Student’s unpaired t-test; ns, no significant; *, p<0.05, ****, p<0.0001.

**Figure 2-figure supplement 2.**
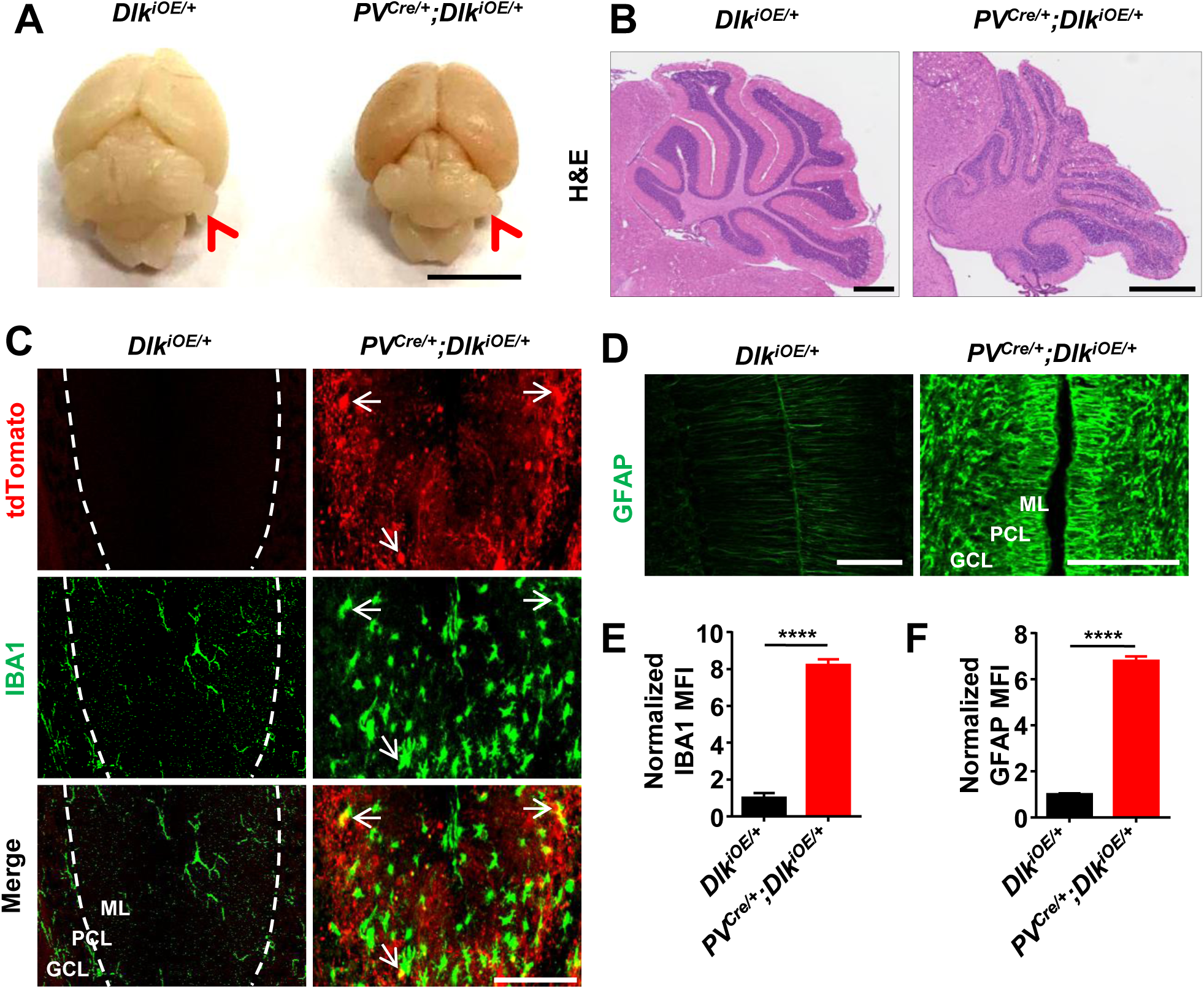
Additional evidence of Purkinje cell degeneration phenotypes caused by DLK activation. A. Representative image of P21 brains of genotypes indicated. Cerebellum (arrowheads) is nearly absent in brain with induced DLK expression in PV^+^ neurons. Scale bar: 5 mm. B. Representative images of hematoxylin and eosin staining of cerebellar sections from P21 mice of genotypes indicated. Scale bars: 500 μm. C. Representative images of cerebellar sections from P21 mice of genotypes indicated, immunostained for IBA1; and tdTomato reporter expression marks Purkinje cells. The arrows show close association between microglia and Purkinje cells. ML: Molecular Layer; PCL: Purkinje Cell Layer; GCL: Granule Cell Layer. Scale bars: 100 μm. D. Representative images of cerebellar sections from P21 mice of genotypes indicated, immunostained for GFAP. E-F. Quantification of normalized IBA1 and GFAP levels in cerebellum of P21 mice. n = 3 per genotype. (E, F). Data shown are means ± SEM. Statistics for E, F: Student’s unpaired t-test; ****, p<0.0001.

**Figure 3-figure supplement 1.**
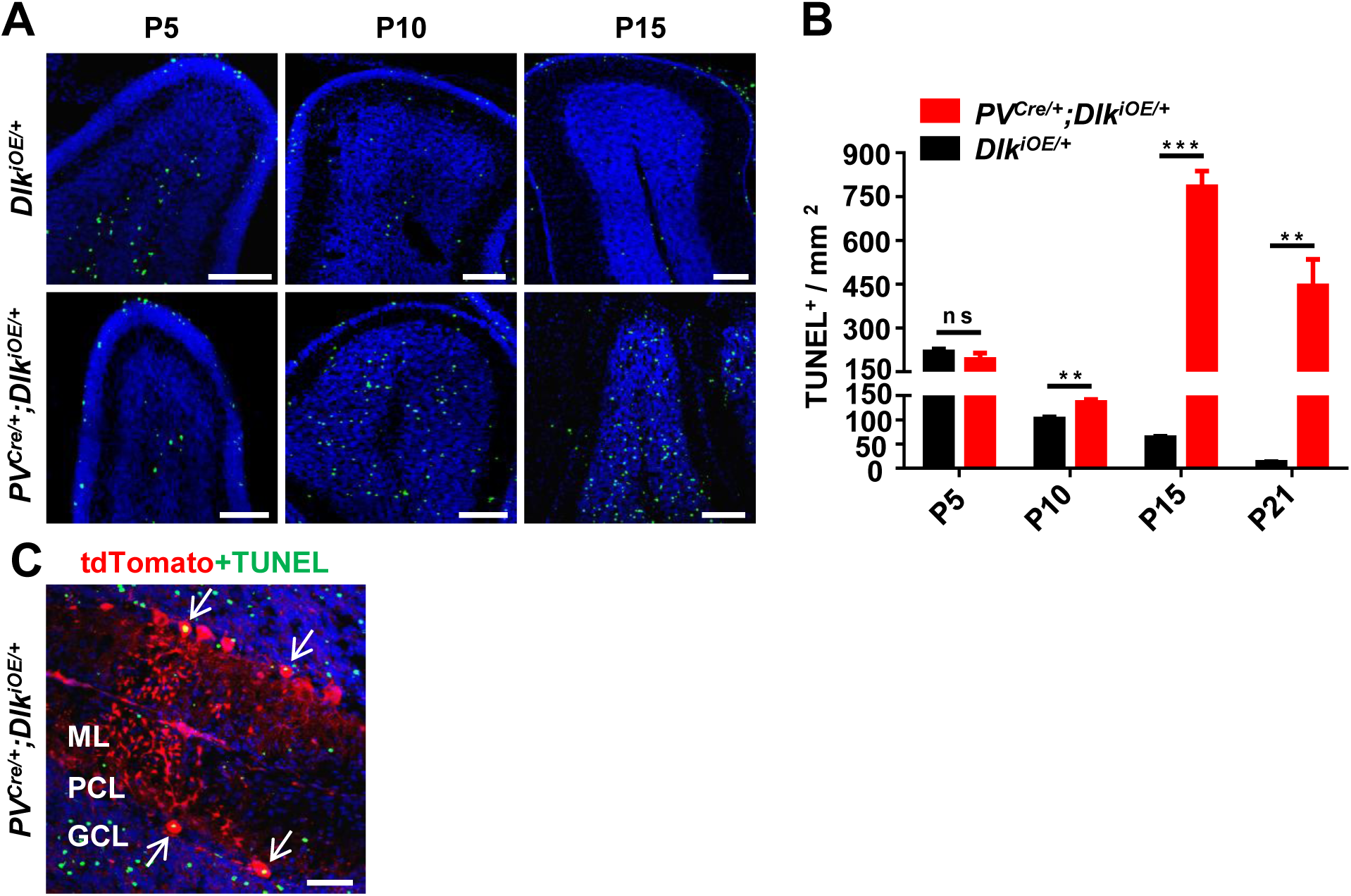
Additional evidence for apoptosis induced by DLK activation. A. Representative images of cerebellar lobules of *Dlk*^*iOE*/+^ and *PV*^*Cre*/+^;*Dlk*^*iOE*/+^ mice from P5 to P15, labeled with TUNEL fluorescence signal (fluorescein-12-dUTP). Scale bars: 100 μm. B. Quantification of TUNEL^+^ cells density in cerebellum. n ≥ 3 per group at each time point. C. Representative image showed the co-localization of TUNEL signal with tdT^+^ Purkinje cells in P15 mice (arrows). ML: Molecular Layer; PCL: Purkinje Cell Layer; GCL: Granule Cell Layer. Scale bar: 50 μm. (B). Data shown are means ± SEM. Statistics for B: Student’s unpaired t-test; ns, no significant; **, p<0.01, ***, p<0.001.

**Figure 3-figure supplement 2.**
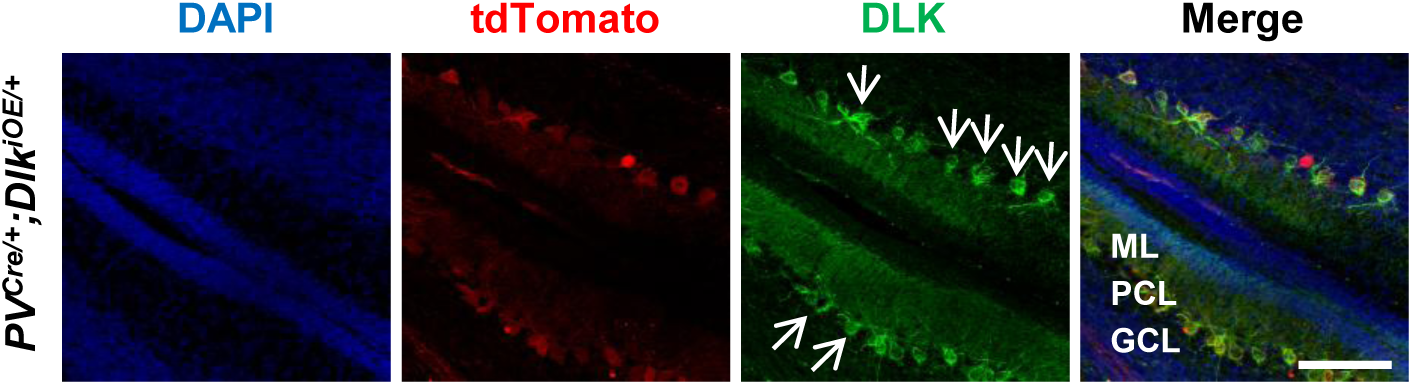
DLK protein localization in Purkinje cells. Representative images of cerebellar section of P10 mice immunostained for DLK; arrows show DLK distribution in Purkinje cells (arrows). ML: Molecular Layer; PCL: Purkinje Cell Layer; GCL: Granule Cell Layer. Scale bar: 100 μm.

**Figure 4-figure supplement 1.**
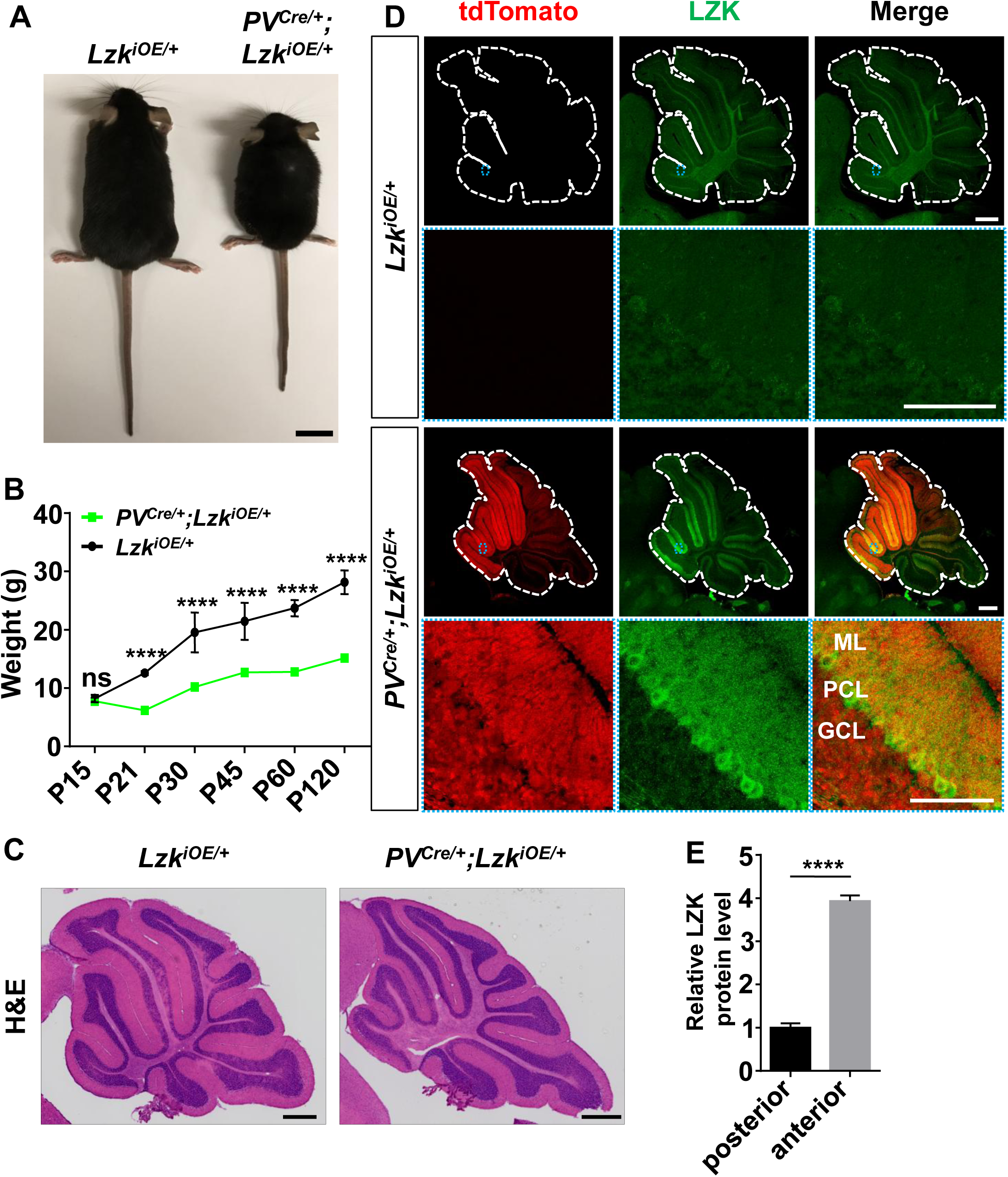
Additional evidence for elevating LZK expression in PV^+^ neurons causing animal growth and movement defects. A. Representative image of P120 mice of genotypes indicated. Scale bar: 2 cm. B. Quantification of the body weight of mice of genotypes indicated from P15 to P120. n ≥ 4 per group at each time point. C. Representative images of hematoxylin and eosin staining of cerebellar sections from P120 mice of genotypes indicated. Scale bars: 500 μm. D. Representative images of Purkinje cells of P21 mice immunostained for LZK. Enlarged blue boxed area show transgenic LZK strongly expressed in dendrites and soma of Purkinje cells in *PV*^*Cre*/+^;*Lzk*^*iOE*/+^ mice. ML: Molecular Layer; PCL: Purkinje Cell Layer; GCL: Granule Cell Layer. Scale bars: 500 μm (whole sections), 100 μm (enlarged views). E. Quantification of relative LZK protein levels in PC soma of anterior and posterior of cerebellum in *PV*^*Cre*/+^;*Lzk*^*iOE*/+^ mice. n = 3. (B, E). Data shown are means ± SEM. Statistics for B, E: Student’s unpaired t-test; ns, no significant; ****, p<0.0001.

**Figure 4-figure supplement 2.**
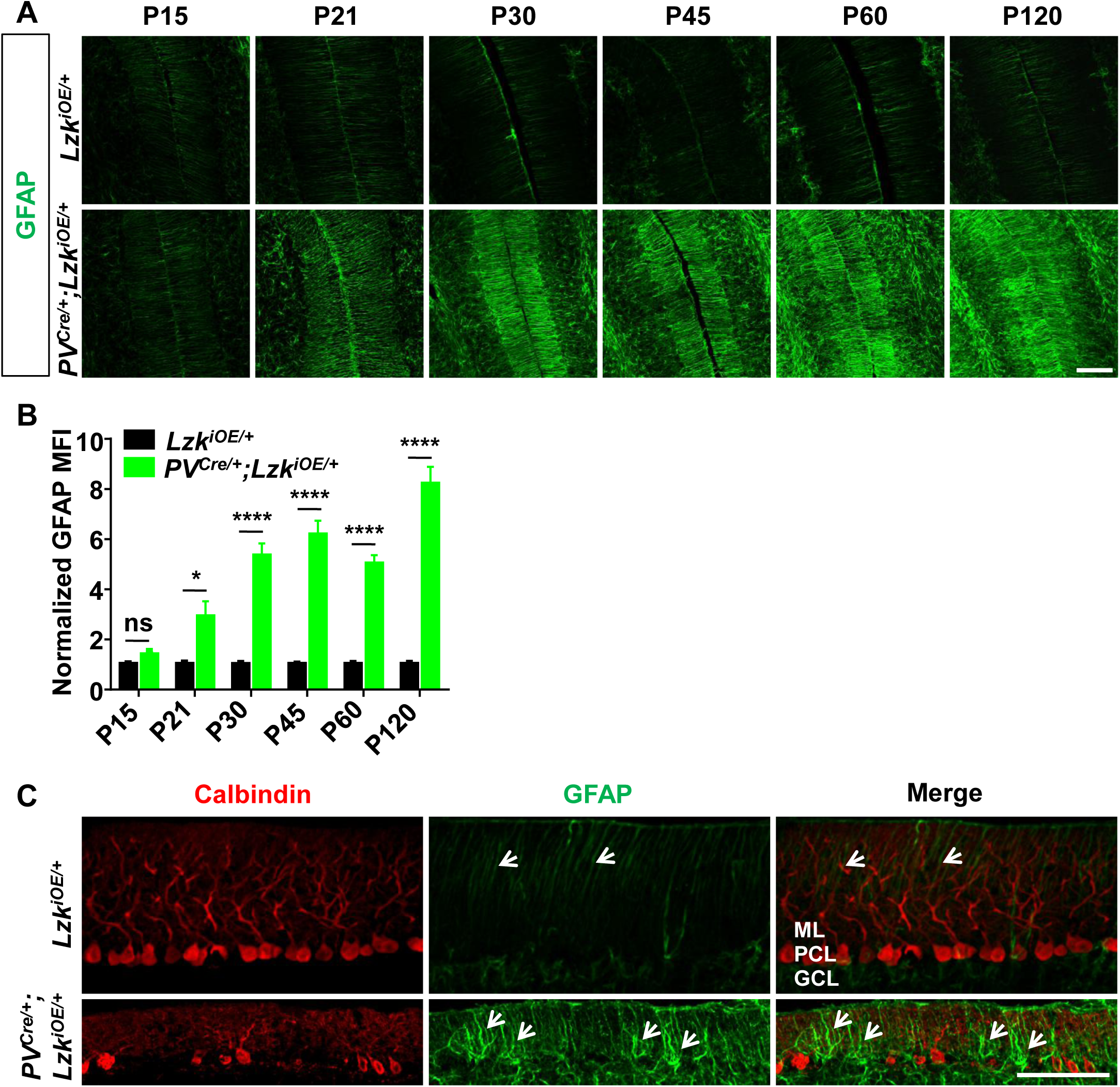
Additional evidence for progressive degeneration of Purkinje cells induced by LZK activation. A. Representative images of GFAP staining of cerebellar astrocytes of the mice from P15 to P120. ML: Molecular Layer; PCL: Purkinje Cell Layer; GCL: Granule Cell Layer. Scale bar: 100 μm. B. Quantification of normalized GFAP levels in cerebellum of the mice from P15 to P120. From P21, mice with induced LZK expression in PV^+^ neurons have significantly higher GFAP levels in cerebellum than control *Lzk*^*iOE*/+^ mice. n = 3 per genotype at each time point. MFI: mean of fluorescence intensity. Data shown are means ± SEM. Statistics: Student’s unpaired t-test; ns, no significant; *, p<0.05, ****, p<0.0001. C. Representative images of Purkinje cells and Bergmann glia in cerebellum of P45 mice, co-immunostained for Calbindin and GFAP. The death of Purkinje cells induced increased reactivity of their adjacent Bergmann glia (arrows). ML: Molecular Layer; PCL: Purkinje Cell Layer; GCL: Granule Cell Layer. Scale bar: 100 μm.

**Figure 5-figure supplement 1.**
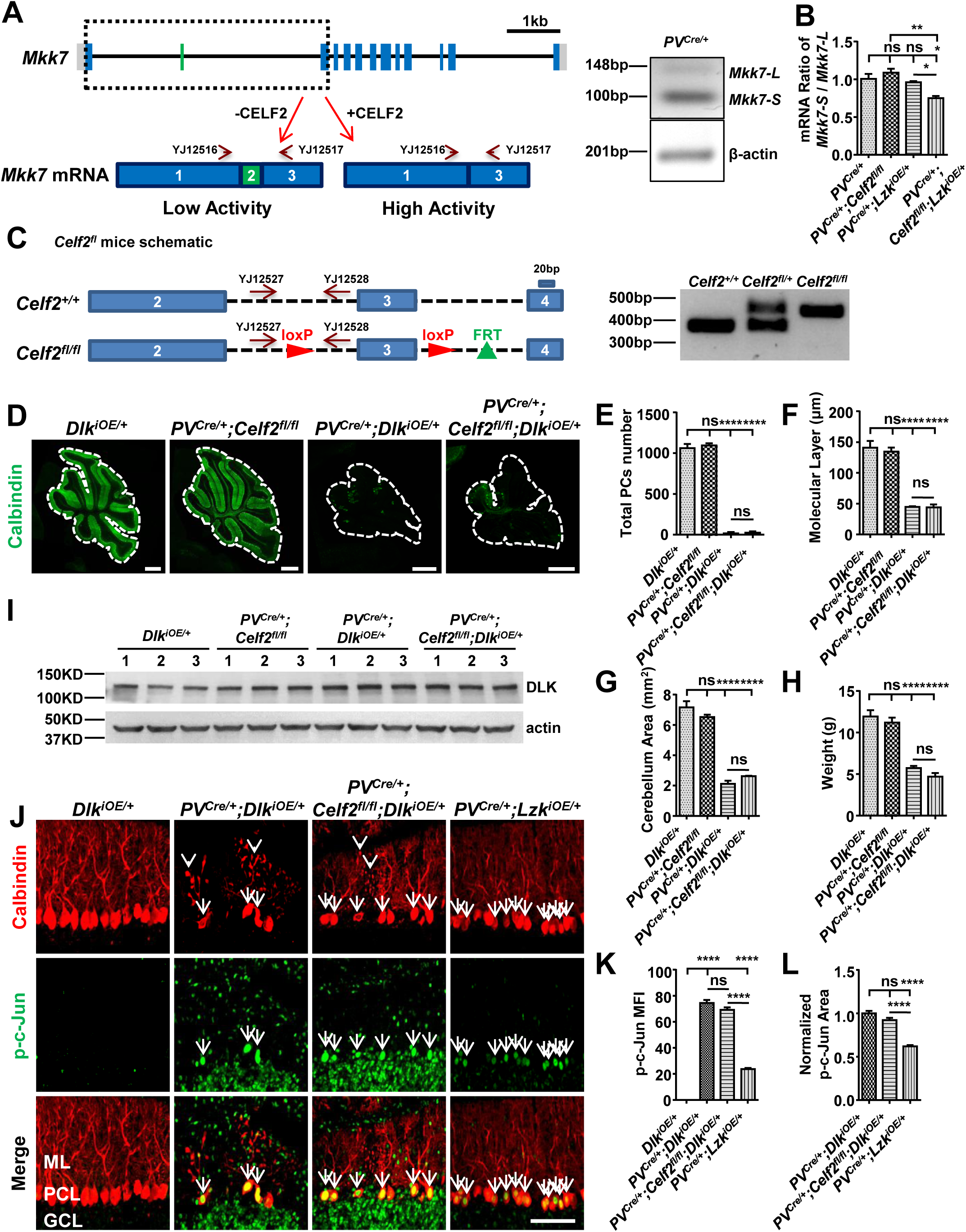
CELF2 is a regulator of *Mkk7* in Purkinje cells. A. Schematic of *mkk7* locus on Chromosome 8, along with primers for mRNA detection of *Mkk7* isoforms. Scale bar: 1kb. CELF2 promotes exclusion of exon 2 of *Mkk7*. The gel image shows the RT-PCR analysis of the *Mkk7* long isoform (*Mkk7-L*) and short isoform (*Mkk7-S*) as well as *β-actin* mRNA levels in cerebellum of *PV*^*Cre*/+^ mouse using primers YJ12516-12519. B. qRT-PCR analyses show the mRNA ratio of *Mkk7-S* / *Mkk7-L* in cerebellum from P21 mice of genotypes indicated. RNA samples are collected from n = 3 mice / genotype. For qRT-PCR, n = 5 replicates / sample. C. The *Celf2*^*fl/fl*^ mice have two loxP sites flanking exon 3, and Cre-mediated excision results in an early stop codon within exon 4, hence a null allele. The homozygotes or heterozygotes of *Celf2*^*fl*^ were identified by PCR using primers YJ12527 and YJ12528. D. Representative images of Calbindin staining of cerebellar sections from P21 mice of genotypes indicated. Scale bars: 500 μm. E. Quantification of total number of PCs in all cerebellar lobules. F. Quantification of the molecular layer thickness in cerebellar lobules V-VI. G. Quantification of the cerebellum area of P21 mice. H. Quantification of the body weight of P21 mice of genotypes indicated. I. The DLK protein levels in cerebellum from P10 mice of genotypes indicated were determined by western blot. n = 3 per group. J. Representative images of Purkinje cells of P15 mice of genotypes indicated, co-immunostained for Calbindin and p-c-Jun. *Celf2* deletion did not reduce the phosphorylation level of c-Jun induced by DLK overexpression in PV^+^ neurons (arrows), nor rescued the dendrite swelling caused by DLK overexpression in PV^+^ neurons (arrowheads). ML: Molecular Layer; PCL: Purkinje Cell Layer; GCL: Granule Cell Layer. Scale bar: 100 μm. K. Quantification of the p-c-Jun levels in Purkinje cells of P15 mice. MFI: mean of fluorescence intensity. L. Quantification of normalized p-c-Jun area in Purkinje cells of P15 mice. DLK overexpression in PV^+^ neurons induces a higher level of p-c-Jun in Purkinje cells than LZK overexpression (arrows). (F-I, K-L). n = 3 per genotype; data shown are means ± SEM. Statistics for B, F-I, K-L: One-way ANOVA; ns, no significant; *, p<0.05, **, p<0.01, ****, p<0.0001.

**Figure 5-figure supplement 2.**
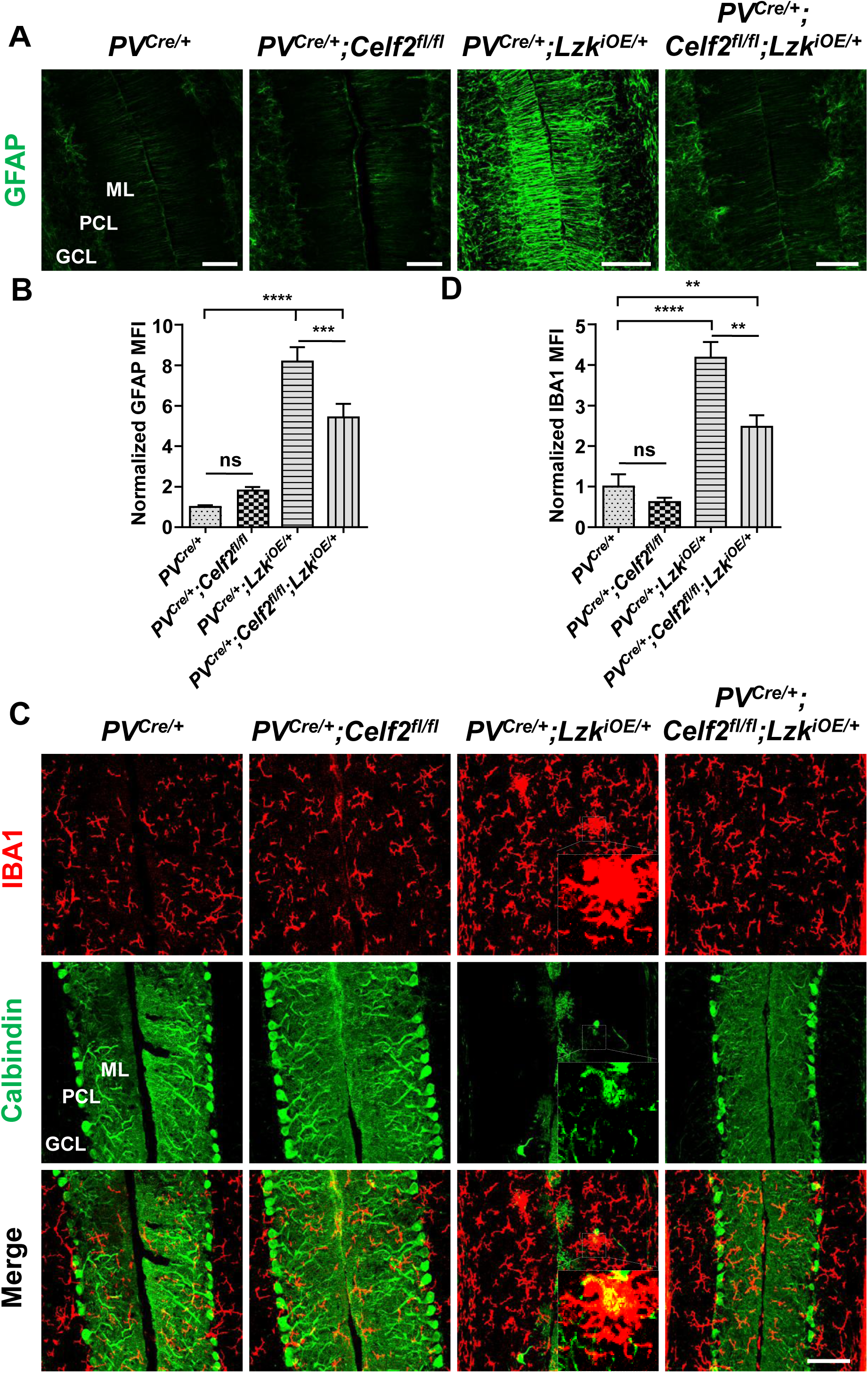
CELF2 deletion reduces astrogliosis and microgliosis associated with LZK activation in PV^+^ neurons. A. Representative images of GFAP staining of cerebellar astrocytes from P120 mice of genotypes indicated. ML: Molecular Layer; PCL: Purkinje Cell Layer; GCL: Granule Cell Layer. Scale bars: 100 μm. B. Quantification of normalized GFAP levels in cerebellum of P120 mice. MFI: mean of fluorescence intensity. C. Representative images of cerebellar sections from P120 mice of genotypes indicated, co-immunostained for IBA1 and Calbindin. Enlarged white boxed areas show dying Purkinje cells caused by LZK activation in PV^+^ neurons were phagocytosed by microglia. Scale bar: 100 μm. ML: Molecular Layer; PCL: Purkinje Cell Layer; GCL: Granule Cell Layer. D. Quantification of normalized IBA1 levels in cerebellum. MFI: mean of fluorescence intensity. (B, D). n = 3 per group; data shown are means ± SEM. Statistics for B, D: One-way ANOVA; ns, no significant; **, p<0.01, ***, p<0.001, ****, p<0.0001.

**Figure 5-figure supplement 3.**
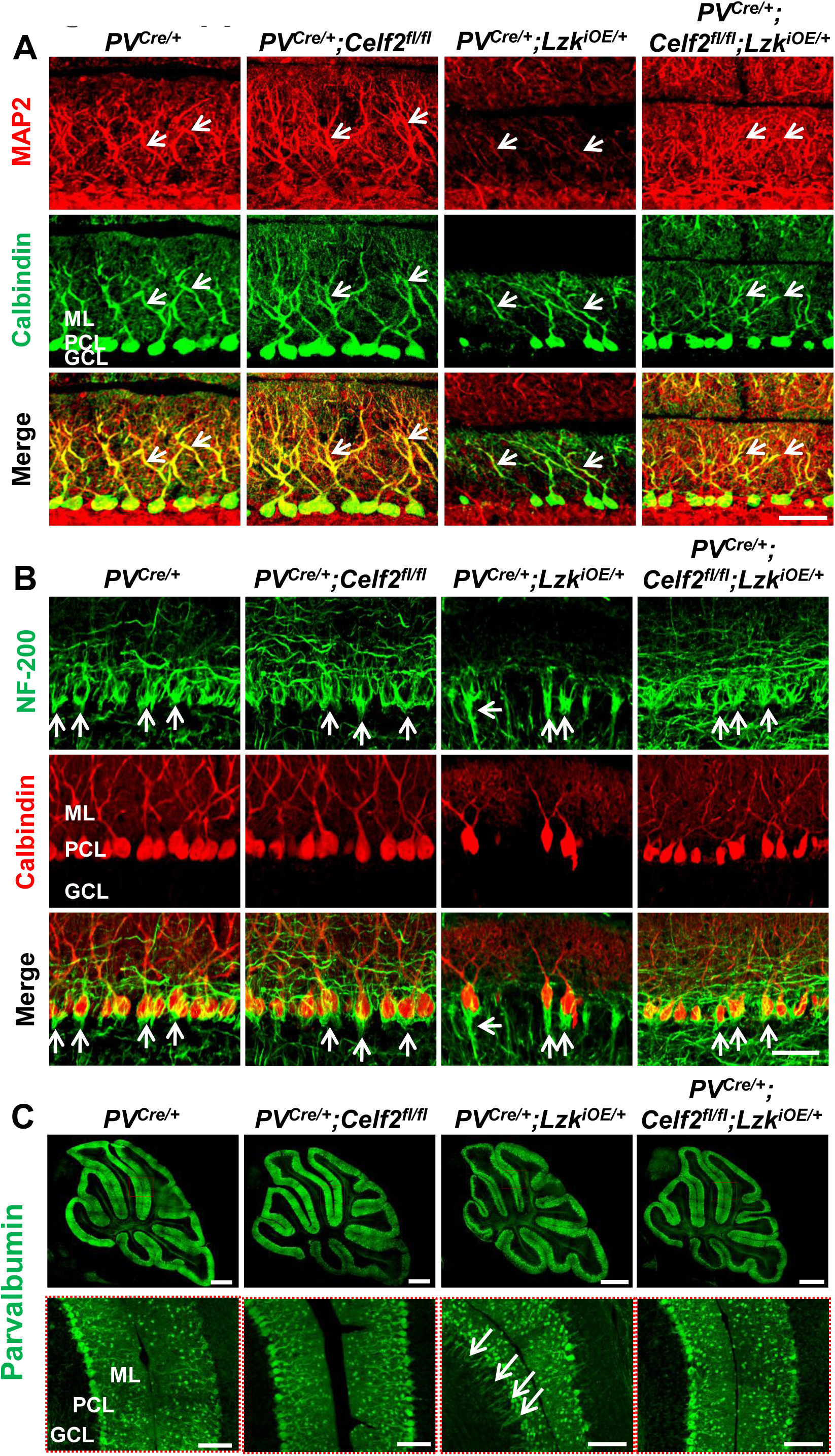
CELF2 deletion rescues reduced expression of MAP2 and NF-200 and pinceau disorganization induced by LZK activation. A. Representative images of cerebellar sections from P120 mice of genotypes indicated, co-immunostained for Calbindin and MAP2. Arrows point MAP2 staining in dendrites of Purkinje cells. B. Representative images of cerebellar sections from P120 mice of genotypes indicated, co-immunostained for NF-200 and Calbindin. LZK overexpression in PV^+^ neurons changes the organization of basket cell pinceaux (arrows) at the initial segment of axons of Purkinje cells, which is suppressed by *Celf2* deletion. (A-B) ML: Molecular Layer; PCL: Purkinje Cell Layer; GCL: Granule Cell Layer. Scale bar: 50 μm. C. Representative images of parvalbumin staining of cerebellar sections from P120 mice of genotypes indicated. Enlarged boxed areas in lower panel show suppression of pinceaux disorganization by *Celf2* deletion. ML: Molecular Layer; PCL: Purkinje Cell Layer; GCL: Granule Cell Layer. Scale bars: 500 μm (upper panel), 100 μm (lower panel).

**Figure 6-figure supplement 1.**
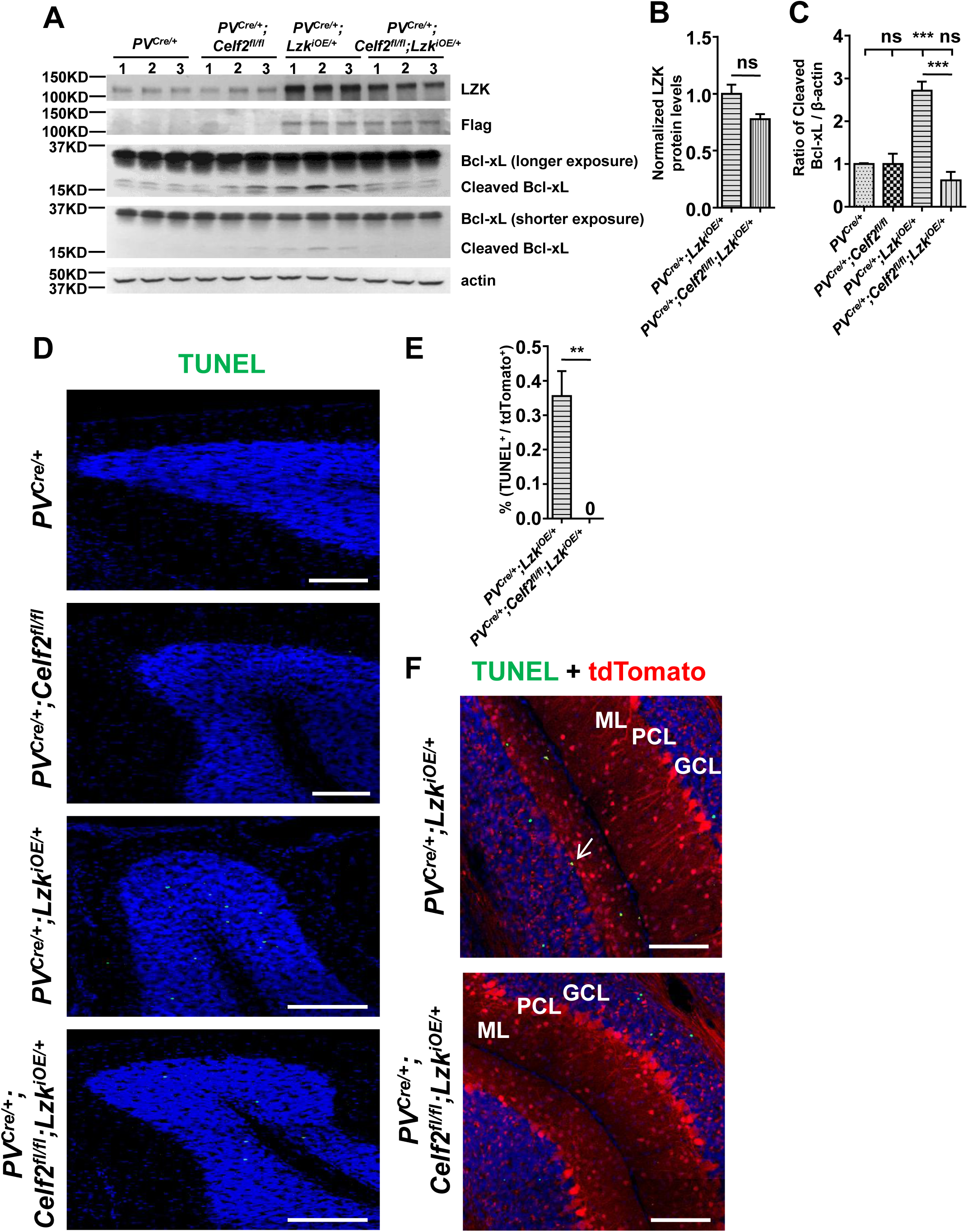
Additional evidence of apoptosis in Purkinje cell degeneration caused by LZK activation. A. Western blot showing LZK and Bcl-xL protein levels in cerebellar extracts from P21 mice of genotypes indicated. The LZK proteins were first immunoprecipitated and then analyzed by western blot. B. Quantification of normalized LZK protein levels. n = 3 per genotype. Data shown are means ± SEM. Statistics: Student’s unpaired t-test; ns, no significant. C. Quantification of the ratio of cleaved Bcl-xL relative to β-actin protein levels. n = 3 per group. Data shown are means ± SEM. Statistics: One-way ANOVA; ns, no significant; ***, p<0.001. D. Representative images of TUNEL fluorescence signal (fluorescein-12-dUTP) in cerebellar lobules of P120 mice of genotypes indicated. Scale bars: 100 μm. E. Quantification of the percentage of TUNEL^+^ Purkinje cells in total Purkinje cells (tdTomato^+^) in *PV*^*Cre*/+^;*Lzk*^*iOE*/+^ and *PV*^*Cre*/+^;*Celf2*^*fl/fl*^;*Lzk*^*iOE*/+^ mice at P120. n = 3 per group. Data shown are means ± SEM. Statistics: Student’s unpaired t-test; **, p<0.01. F. Representative images of cerebellar sections of *PV*^*Cre*/+^;*Lzk*^*iOE*/+^ and *PV*^*Cre*/+^;*Celf2*^*fl/fl*^;*Lzk*^*iOE*/+^ mice labeled with TUNEL fluorescence signal and reporter tdTomato. The TUNEL signal was co-localized with Purkinje cells in *PV*^*Cre*/+^;*Lzk*^*iOE*/+^ mice (arrow), but not in *PV*^*Cre*/+^;*Celf2*^*fl/fl*^;*Lzk*^*iOE*/+^ mice. ML: Molecular Layer; PCL: Purkinje Cell Layer; GCL: Granule Cell Layer. Scale bars: 100 μm.

**Figure 7-figure supplement 1.**
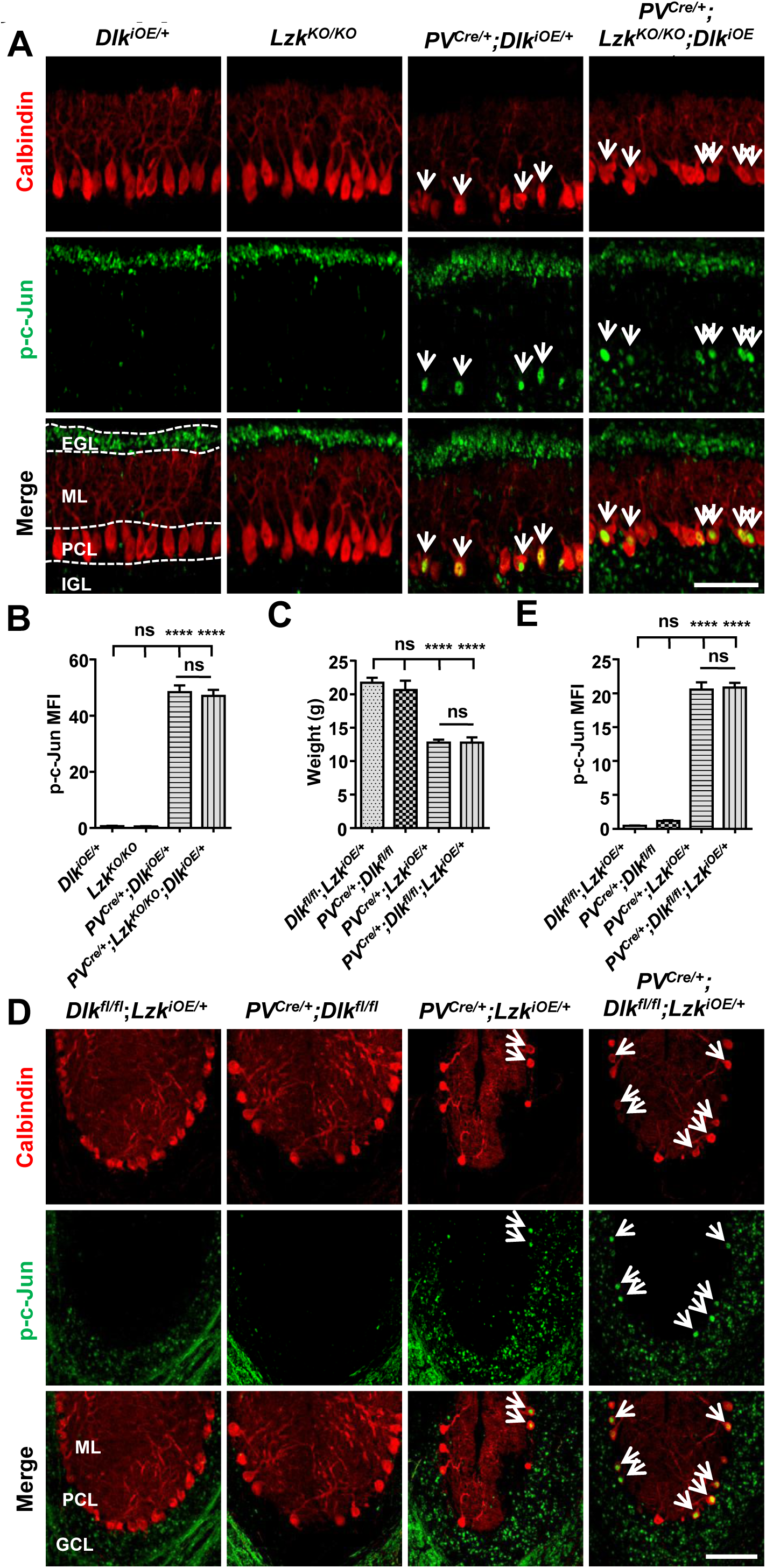
Levels of p-c-Jun induced by activation of DLK or LZK is not affected by loss of *Lzk* or *Dlk*, respectively. A. LZK deletion did not attenuate the phosphorylation of c-Jun induced by DLK overexpression in PV^+^ neurons. Shown are representative images of cerebellar sections of P10 mice, co-immunostained for p-c-Jun and Calbindin. EGL: External Granule Layer; ML: Molecular Layer; PCL: Purkinje Cell Layer; IGL: Internal Granule Layer. Scale bar: 100 μm. B. Quantification of the p-c-Jun levels in Purkinje cells of P10 mice of genotype indicated. MFI: mean of fluorescence intensity. n = 3 per genotype. C. Quantification of the body weight of P60 mice of genotypes indicated. n ≥ 3 per genotype. D. DLK deletion did not attenuate the phosphorylation of c-Jun induced by LZK overexpression in PV^+^ neurons. Shown are representative images of cerebellar lobules of P60 mice, co-immunostained for Calbindin and p-c-Jun. ML: Molecular Layer; PCL: Purkinje Cell Layer; GCL: Granule Cell Layer. Scale bar: 100 μm. E. Quantification of the p-c-Jun levels in Purkinje cells of P60 mice. MFI: mean of fluorescence intensity. n = 3 per group. (B, C, E). Data shown are means ± SEM. Statistics for B, C, E: One-way ANOVA; ns, no significant; ****, p<0.0001.

## Supplementary Videos

**Video 1. Locomotor deficits in P15 *PV***^***Cre*/+**^**;*Dlk***^***iOE*/+**^ **mice**.

A representative P15 old *PV*^*Cre*/+^;*Dlk*^*iOE*/+^ mouse exhibited difficulty moving forward and drags its abdomen along the ground, as well as tremors.

**Video 2. Locomotor deficits in P120 *PV***^***Cre*/+**^**;*Lzk***^***iOE*/+**^ **mice**.

A representative P120 old *PV*^*Cre*/+^;*Lzk*^*iOE*/+^ mouse exhibited severe limp while walking and tremors.

